# Network-based investigation of petroleum hydrocarbons-induced ecotoxicological effects and their risk assessment

**DOI:** 10.1101/2024.07.18.604159

**Authors:** Ajaya Kumar Sahoo, Shreyes Rajan Madgaonkar, Nikhil Chivukula, Panneerselvam Karthikeyan, Kundhanathan Ramesh, Shambanagouda Rudragouda Marigoudar, Krishna Venkatarama Sharma, Areejit Samal

**Affiliations:** The Institute of Mathematical Sciences (IMSc), Chennai, India; Homi Bhabha National Institute (HBNI), Mumbai, India; National Centre for Coastal Research, Ministry of Earth Sciences, Government of India, Pallikaranai, Chennai, India

**Author notes:** A.K. Sahoo and S.R. Madgaonkar contributed equally to this work and should be considered as Joint-First authors. Corresponding author (A. Samal) **Address for correspondence**: Areejit Samal Computational Biology Group, The Institute of Mathematical Sciences (IMSc), CIT Campus, Taramani, Chennai 600113 India.

**Keywords:** Petroleum hydrocarbons, Adverse outcome pathway, Stressor-AOP network, Stressor-species network, Ecological risk assessment, Species sensitivity distribution

## Abstract

Petroleum hydrocarbons (PHs) are compounds composed mostly of carbon and hydrogen, originating from crude oil and its derivatives. PHs are primarily released into the environment through the diffusion of oils, resulting from anthropogenic activities like transportation and offshore drilling, and accidental incidents such as oil spills. Once released, these PHs can persist in different ecosystems and cause long-term detrimental ecological impacts. While the hazards associated with such PH contaminations are often assessed by the concentrations of total petroleum hydrocarbons in the environment, studies focusing on the risks associated with individual PHs are limited. Here, we leveraged different network-based frameworks to explore and understand the adverse ecological effects associated with PH exposure. First, we systematically curated a list of 320 PHs from published reports. Next, we integrated biological endpoint data from toxicological databases, and constructed a stressor-centric adverse outcome pathway (AOP) network linking 75 PHs with 177 ecotoxicologically-relevant high confidence AOPs within AOP-Wiki. Further, we relied on stressor-species network constructions, based on reported toxicity concentrations and bioconcentration factors data for 80 PHs and 28 PHs, respectively, and found that crustaceans are documented to be affected by many of these PHs. Finally, we utilized the aquatic toxicity data within ECOTOX to construct species sensitivity distributions for polycyclic aromatic hydrocarbons (PAHs) prioritized by the US EPA, and derived their corresponding hazard concentrations (HC05) that protect 95% of species in the aquatic ecosystem. Overall, this study highlights the importance of using network-based approaches and risk assessment methods to understand the PH-induced toxicities effectively.

## 1. Introduction

Petroleum hydrocarbons (PHs) are compounds consisting mostly of carbon and hydrogen besides other elements that originate from crude oil and its derivatives such as gasoline and diesel, among others (Kuppusamy et al., 2020). PHs are released into the environment primarily through the diffusion of oils, stemming from anthropogenic activities such as transportation and offshore drilling, as well as accidental incidents like oil spills (Tornero and Hanke, 2016; Zheng and Richardson, 1999). Eventually, PHs are absorbed through various exposure routes, including ingestion, dermal contact, and inhalation, where they can bioaccumulate and lead to carcinogenic, developmental, and endocrine toxicities in humans and other species (Almeda et al., 2013b, 2013a; Pritsos et al., 2017; Takeshita et al., 2021). PHs can be broadly classified into aliphatic and aromatic hydrocarbons, with the aromatic PHs being widely studied due to their higher stability, water solubility, and environmental persistence (Alford et al., 2014; Al-Hawash et al., 2018). Moreover, the United States Environmental Protection Agency (US EPA) has designated 16 of these polycyclic aromatic hydrocarbons (PAHs) as priority pollutants based on their environmental prevalence and persistence (Hussar et al., 2012; Keith, 2015; Lawal, 2017). Therefore, identifying PHs and understanding their adverse effects will enable the formulation of effective mitigation and remediation strategies for PH contamination.

Accidental oil spillage can contribute to increased environmental PH concentrations causing detrimental impact to diverse ecological habitats. The Deepwater Horizon oil spill in the northern Gulf of Mexico, off the coast of Louisiana, USA (Beyer et al., 2016), is one of the largest oil spills documented to have long-term ecological impacts on fishes, deep ocean corals and oysters, and reduction in the population of marine mammals, sea turtles and seabirds (Barron et al., 2020; Pasparakis et al., 2019). Similarly, the oil spill in the coastal waters of Ennore near Chennai, India, has been documented to reduce water quality and impact marine biota including phytoplankton, zooplankton, benthic communities and vertebrates like fishes and sea turtles (Begum et al., 2022). Although the hazards associated with such oil contaminations are often assessed by measuring total petroleum hydrocarbon (TPH) concentrations in the affected environments (Sammarco et al., 2013; Sivagami et al., 2019), there is a lack of studies focusing on the risks posed by individual PHs.

With the rise in technological advancements and the call for new approach methodologies (National Research Council, 2007), *in silico* tools that integrate heterogenous toxicological data have emerged as effective frameworks for evaluating the ecotoxicological risks of chemicals (Scholz et al., 2013; Thomas et al., 2019). In line with this, the Adverse Outcome Pathway (AOP) framework was developed to provide an effective translation of mechanistic data across different levels of biological organization to endpoints relevant to ecological risk assessment (Ankley et al., 2010; Baudiffier et al., 2024; Jeong and Choi, 2019; Kramer et al., 2011; Leist et al., 2017). The AOP framework organizes stressor-induced biological events, termed as key events (KEs), in a sequential setup where the KEs are linked through causal relationships called key event relationships (KERs) (Villeneuve et al., 2014a, 2014b). Networks built from these AOPs, after integrating various stressor-specific information, have proven to provide a comprehensive understanding of the diverse mechanisms underlying stressor-induced toxicities (Chai et al., 2021; del Giudice et al., 2024; Knapen et al., 2018; Sahoo et al., 2024b, 2024a; Villeneuve et al., 2018). ECOTOX is the most comprehensive knowledgebase providing toxicity data for individual chemicals across diverse aquatic and terrestrial species (Olker et al., 2022). As one of the largest ecotoxicological databases, its integration with the AOP framework will aid in understanding the complex ecotoxicological mechanisms underlying stressor-induced effects through AOP networks (Fay et al., 2017; Russom et al., 2014). Additionally, stressor-species networks constructed using diverse toxicological information from ECOTOX have facilitated the investigation of interactions between stressors and various ecological species (Wang et al., 2024). Moreover, the toxicity data available in ECOTOX has enabled the construction of species sensitivity distributions (SSDs), which help elucidate the interspecies variability in response to environmental contaminants (Karthikeyan et al., 2021; Posthuma et al., 2019; Tian et al., 2020). Importantly, the constructed SSDs can aid in ecological risk assessment by providing threshold concentrations of chemicals that are protective of a large proportion of species in a given environment (Dowse et al., 2013; EPA, 1998; Posthuma et al., 2001). Therefore, *in silico* approaches that integrate ecotoxicological data can aid in the ecological risk assessment of individual PHs.

In this study, we curated a list of PHs from the reports published by the Total Petroleum Hydrocarbon Criteria Working Group (TPHCWG) (Gustafson et al., 1997; Potter and Simmons, 1998) on the presence of PHs across various fuel oils, including crude oil, and explored their toxicities by leveraging the ecotoxicological data available in ECOTOX (Olker et al., 2022). First, we curated ecotoxicologically-relevant high confidence AOPs from AOP-Wiki (https://aopwiki.org) based on their taxonomic applicability annotations. Then, we systematically integrated biological endpoint data corresponding to these PHs from three sources namely, ToxCast (Dix et al., 2007), Comparative Toxicogenomics Database (CTD) (Davis et al., 2023) (https://ctdbase.org/) and ECOTOX to identify associated KEs from AOP-Wiki, and subsequently constructed a stressor-AOP network for PHs. We utilized this constructed stressor-AOP network to identify highly relevant AOPs associated with benzo[a]pyrene-induced toxicities, and performed a case study to understand the rationale behind its ecotoxicological effects. Further, we utilized the toxicity concentrations and bioconcentration factors data within ECOTOX to construct stressor-species networks, and leveraged them to understand the ecotoxicological effects of PHs across diverse species. Finally, we utilized the acute toxicity data available in ECOTOX to compute the SSDs for the EPA priority PAHs, and derived their hazard concentrations for aquatic environments. In sum, this study systematically identifies PHs from published reports, and conducts network-based investigation and ecological risk assessments to address their ecotoxicological effects.

## 2. Materials and methods

### 2.1. Compilation and curation of petroleum hydrocarbons

Organic compounds originating from crude oil, and consisting mostly carbon and hydrogen atoms are termed as petroleum hydrocarbons (PHs) (Kuppusamy et al., 2020). Total petroleum hydrocarbons (TPHs) refer to the total recoverable concentrations of PHs measured in an environmental sample (Kuppusamy et al., 2020). The Total Petroleum Hydrocarbon Criteria Working Group (TPHCWG), consisting of representatives from industry, academia, and government, had convened to provide extensive technical information relevant for risk assessment of PHs in petroleum contaminated sites (Weisman, 1998). TPHCWG compiled their findings and data into a series of reports dealing with analytical methods for quantifying TPH, composition of various fuel mixtures, fate and transport of TPH, fraction-based reference dose and reference concentration development, and framework for human health risk-based evaluation of PH contaminated sites.

In this study, we relied on TPHCWG report Volume 2 (Potter and Simmons, 1998), titled ‘Composition of Petroleum Mixtures,’ and TPHCWG report Volume 3 (Gustafson et al., 1997), titled ‘Selection of Representative TPH Fractions Based on Fate and Transport Considerations’, to retrieve PHs present in 11 petroleum mixtures namely, gasoline, diesel, kerosene, number 2 fuel oil, number 6 fuel oil, JP-4, JP-5, JP-7, JP-8, lubricating and motor oils, and crude oil. We mapped these retrieved chemicals to standardized chemical information from PubChem (https://pubchem.ncbi.nlm.nih.gov) and Chemical Abstracts Service (CAS) (https://commonchemistry.cas.org/), and subsequently compiled a list of 320 unique PHs (Table S1). The workflow for identifying these 320 PHs is described in Figure 1a. Finally, for each of these identified PHs, we obtained their corresponding chemical structures from PubChem, and employed RDKit (https://www.rdkit.org) to classify them as aliphatic, monocyclic aromatic, or polycyclic aromatic hydrocarbons (Table S1). Further, we employed ClassyFire (Djoumbou Feunang et al., 2016) (http://classyfire.wishartlab.com) to obtain additional chemical categorizations, namely, chemical Kingdom, Superclass and Class (Table S1).

**Figure 1:**
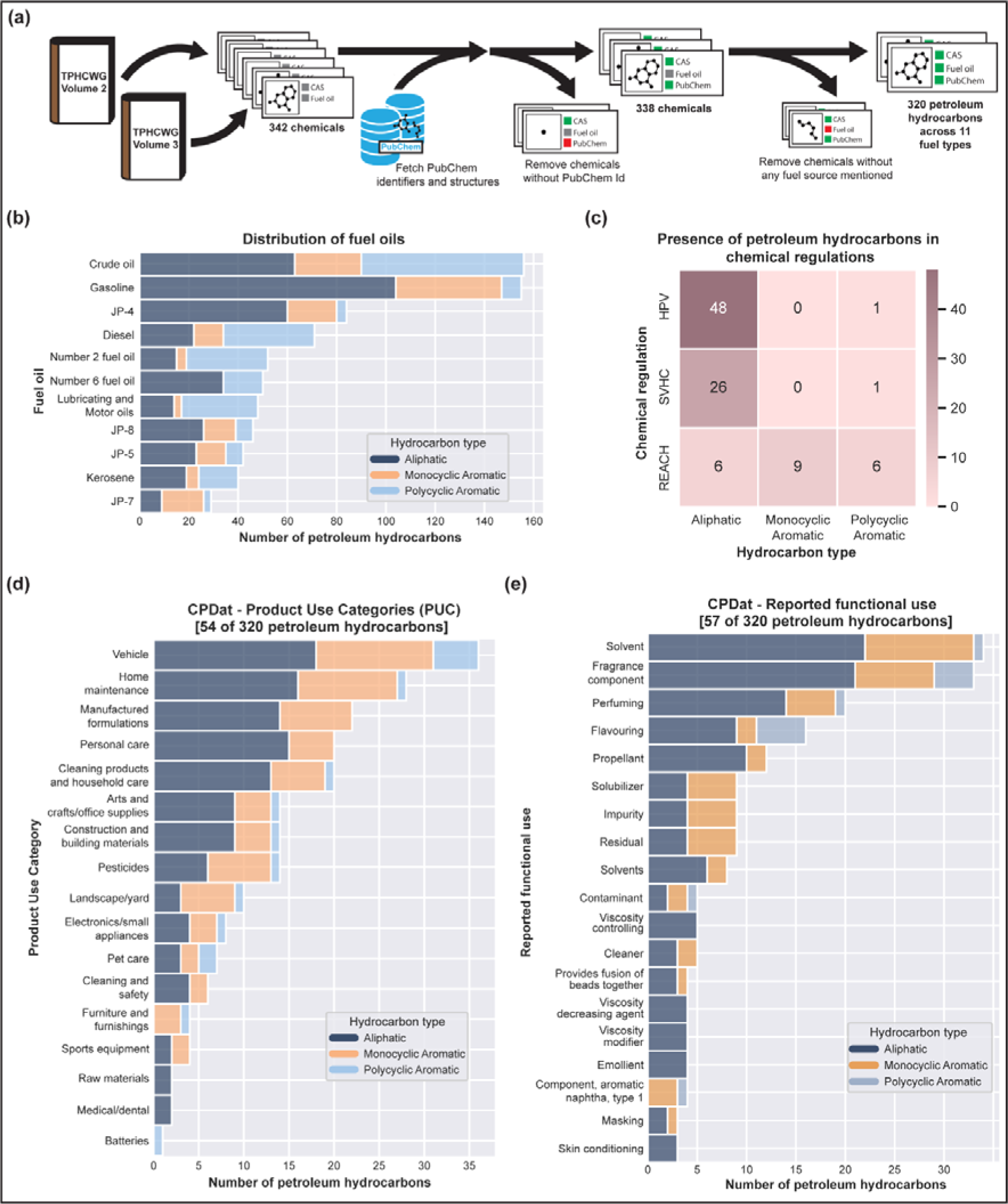
Curation and exploration of petroleum hydrocarbons (PHs). **(a)** Workflow followed to curate a list of 320 PHs from published literature. **(b)** Distribution of different types of PHs across various fuel oils. **(c)** Heatmap depicting the presence of PHs in chemical regulations. The number of different types of PHs in each of the chemical regulations is denoted in the heatmap. **(d)** Distribution of different types of PHs across various product use categories from CPDat. **(e)** Distribution of different types of PHs across various functional uses reported in CPDat which are associated with at least three PHs.

### 2.2. Identification of ecotoxicologically-relevant ‘high confidence’ AOPs within AOP-Wiki

Adverse outcome pathway (AOP) is a modular toxicological framework that captures the different biological events (termed as Key Events or KEs) underlying the process that originates at a biological target that is perturbed by a stressor (termed as Molecular Initiating Event or MIE), and terminates at an Adverse Outcome (AO) of regulatory relevance (Ankley et al., 2010). In an AOP, successive KEs are connected through a directed relationship, termed as Key Event Relationship (KER). AOP-Wiki (https://aopwiki.org/) is the largest publicly accessible global repository that systematically catalogs all the developed AOPs till date, including the scientific evidence supporting these AOPs at various levels. Therefore, we relied on AOP-Wiki to retrieve the AOPs.

First, we downloaded the XML file (released on 1 January 2024) from ‘Project Downloads’ page in AOP-Wiki, and using an in-house python script, we extracted various information associated with the AOPs like their title, identifier, associated KEs (including MIEs and AOs), KERs, stressors, Organisation for Economic Co-operation and Development (OECD) and Society for the Advancement of Adverse Outcome Pathways (SAAOP) status, biological applicability information such as taxonomy, sex, life-stage of the organism, and the weight of evidence. The AOP documentation within AOP-Wiki is constantly updated based on novel understanding and experimental information as and when they are available, and thus are considered as living documents (https://aopwiki.org/handbooks/4). Therefore, to assess the quality and completeness of the AOP data available within AOP-Wiki, we followed a systematic workflow developed in our previous work (Figure S1) (Sahoo et al., 2024b).

We first manually checked and removed ‘archived’ AOPs, AOPs that lacked any KEs or KERs, and AOPs comprising undefined KEs. Then, we employed NetworkX (Hagberg et al., 2008) to identify the directed paths between MIE(s) and AO(s) within each AOP, and removed AOPs if they were disconnected. Finally, through this extensive manual and computational effort, we identified 328 non-empty, connected, complete and high quality AOPs, which we designate as ‘high confidence’ AOPs.

Next, we followed the workflow proposed by Jagiello *et al*. (Jagiello et al., 2022) to identify AOPs relevant for ecotoxicology based on their associated taxonomic applicability. We first filtered out the high confidence AOPs that lacked any taxonomic applicability information. Then, we filtered out the AOPs if their taxonomic applicability contained only terms related to humans. Through this systematic process, we identified 195 of the 328 high confidence AOPs to be relevant for ecotoxicology, which we designate as ‘ecotoxicologically-relevant AOPs’.

Table S2 contains the list of 195 ecotoxicologically-relevant AOPs obtained through our systematic workflow. These 195 ecotoxicologically-relevant AOPs comprise 727 unique KEs (Table S3) and 1047 unique KERs (Table S4).

### 2.3. Identification of ecotoxicologically-relevant KEs associated with petroleum hydrocarbons

In this study, we aimed to analyze the ecotoxicity of the PHs through the AOP framework. To achieve this, we first identified the ecotoxicologically-relevant KEs associated with PHs from three different sources namely, ToxCast (Dix et al., 2007), Comparative Toxicogenomics Database (CTD) (Davis et al., 2023) (https://ctdbase.org/), and ECOTOX (Olker et al., 2022). Here, we note that AOP-Wiki did not catalog any PH as a prototypical stressor, and therefore we did not rely on AOP-Wiki to obtain KEs associated with PHs.

#### 2.3.1. Using ToxCast

The US EPA’s chemical prioritization program, ToxCast, provides several high-throughput screening assay data points to assess the toxicity of several thousand environmental chemicals (Dix et al., 2007). Based on our previous work (Sahoo et al., 2024b, 2024a), we followed a systematic pipeline to identify the KEs associated with the curated PHs by relying on their ToxCast assay endpoints.

Briefly, we accessed the ToxCast invitrodb version 4.1 dataset and identified active assay endpoints (‘hitc’ ≥ 0.9) (Feshuk et al., 2023) associated with PHs from the ‘mc5-6_winning_model_fits-flags_invitrodb_v4_1_SEPT2023.csv’ file. We additionally identified the ‘activatory’ or ‘inhibitory’ response of the PHs based on the ‘top’ value of the corresponding winning model from the ‘mc4_all_model_fits_invitrodb_v4_1_SEPT2023.csv’ file (Feshuk et al., 2023). Next, we relied on the ‘cytotox_invitrodb_v4_1_SEPT2023.xlsx’ file to identify and discard the cytotoxicity-associated bursts associated with PHs, as they result from non-specific reporter gene activations associated with cell stress and cytotoxicity (Feshuk et al., 2023; Judson et al., 2016; Sahoo et al., 2024a).

Finally, to identify the ecotoxicologically-relevant ToxCast assay endpoints, we relied on the ‘assay_annotations_invitrodb_v4_1_SEPT2023.xlsx’ file to obtain the organism information associated with the assay endpoints. We removed chemical-assay endpoint pairs related to humans. Through this effort, we identified 390 ecotoxicologically-relevant ToxCast assay endpoints which are associated with 56 PHs.

Next, we relied on the assay annotations for these ToxCast endpoints to manually inspect and assess endpoints that can be potentially linked to KEs. Through this process, we identified 14 KEs from ecotoxicologically-relevant AOPs which are associated with 19 assay endpoints (Table S5). Briefly, ToxCast provides the assay endpoints, associated genes, and ‘activatory’ or ‘inhibitory’ information for every tested chemical. We leveraged this information and manually mapped it to the KEs within AOP-Wiki based on their title, object identifier and object name (Sahoo et al., 2024b).

#### 2.3.2. Using CTD

Comparative Toxicogenomics Database (CTD) (Davis et al., 2023) (https://ctdbase.org/) compiles data on chemical-gene/protein, chemical-phenotype, chemical-disease and gene-disease associations (or links) from published literature. Based on our previous work (Sahoo et al., 2024b, 2024a), we utilized chemical (C), gene (G), phenotype (P), and disease (D) tetramers (CGPD-tetramers) within CTD data to identify the ecotoxicologically-relevant KEs associated with the curated PHs. To identify the gene-phenotype associations, we relied on the GO term annotations from the NCBI Gene resource (https://ftp.ncbi.nih.gov/gene) (last accessed on 22 April 2024).

We downloaded the March 2024 release of the CTD data and retrieved CGPD-tetramers associated with the curated PHs. Here, we observed that CTD additionally provides information on the organisms associated with every chemical-gene and chemical-phenotype link. Therefore, to identify ecotoxicologically-relevant CGPD-tetramers, we removed CGPD-tetramers where both the chemical-gene and chemical-phenotype links were related to human. Through this process, we identified 8454 ecotoxicologically-relevant CGPD-tetramers comprising 18 PHs, 904 genes, 210 phenotypes, and 148 diseases (Table S6). Additionally, we employed the GOSim package (Fröhlich et al., 2007) in R programming language to identify the neighboring GO terms of phenotypes. Finally, we manually inspected phenotypes (including their neighbor terms) and diseases, and identified that 90 KEs are associated with 49 phenotypes and 35 KEs are associated with 54 diseases (Table S5). The mapped phenotypes were associated with 18 PHs while the mapped diseases were associated with 15 PHs.

#### 2.3.3. Using ECOTOX

US EPA’s ECOTOX (Olker et al., 2022) is one of the largest ecotoxicological knowledgebases that has compiled manually curated ecotoxicology information for more than 12,000 chemicals across more than 13,000 terrestrial and aquatic species. Importantly, ECOTOX relies on a standardized pipeline to extract various toxicity data from published literature, including the biological effects observed in organisms after exposure to environmental chemicals. Therefore, we utilized the ECOTOX dataset to identify the KEs associated with the curated PHs.

First, we downloaded the latest ECOTOX dataset (released in March 2024) by selecting the ‘Download ASCII Data’ option on the ECOTOX website (https://cfpub.epa.gov/ecotox/). Then, we used an in-house python script to parse and extract data associated with the curated PHs from ‘tests.txt’, ‘results.txt’, ‘chemicals.txt’ and ‘species.txt’ files. We observed that ECOTOX provides curated data points such as biological effects associated with the chemical (Effect), the parameter that measures the corresponding biological effect (Measurement) and the trend (Trend) of this parameter with respect to a control. Here, we removed data points where the Measurement value is ‘Not Reported’ or is empty, as they lack information on the adverse effects caused by such chemicals. Finally, we manually inspected the Effect, Measurement and Trend parameters, and identified that 101 KEs are associated with 74 PHs (Table S5).

Overall, we identified 206 ecotoxicologically-relevant KEs associated with 75 out of the 320 PHs in our curated list through biological endpoints data from three sources: ToxCast, CTD and ECOTOX.

### 2.4. Construction and visualization of stressor-AOP network for petroleum hydrocarbons

Stressor-AOP network can capture a diverse array of AOPs linked to stressors, and thus, provide a comprehensive understanding of the adverse effects induced by a chemical. To understand the adverse effects induced by the PHs, we constructed a bipartite graph between the PHs and the ecotoxicologically-relevant AOPs, where PHs are linked to an AOP through an associated KE. Further, we characterized the stressor-AOP links between PHs and AOPs by computing the coverage score and level of relevance. The coverage score of an AOP is obtained by taking the ratio of the number of KEs which are associated with PHs to the total number of KEs in that AOP. Further, the level of relevance is assessed based on the following five-level criterion:

● Level 1: stressor is associated with the KE of the AOP but not associated with any MIE or AO of the AOP
● Level 2: stressor is associated with the AO of the AOP but not associated with any MIE of the AOP
● Level 3: stressor is associated with the MIE of the AOP but not associated with any AO of the AOP
● Level 4: stressor is associated with both MIE and AO of the AOP
● Level 5: stressor is associated with both MIE and AO of the AOP and there exist a directed path between the associated MIE and AO

We visualized the stressor-AOP network using Cytoscape (Shannon et al., 2003), where the stressor-AOP link is annotated by the coverage score and the level of relevance. Table S7 provides the complete stressor-AOP network for the PHs.

### 2.5. Construction and visualization of stressor-species network using ECOTOX data

For a chemical, ECOTOX provides toxicity concentration value at which the biological effect is observed and the bioconcentration factor, across different species. To construct a stressor-species network using the toxicity concentration value, we first retrieved the chemical concentrations for acute toxicity endpoints (LC_50_ and EC_50_) for the curated list of 320 PHs across different species. Note that, ECOTOX provides the toxicity concentration values in different units of measurement, therefore we standardized these values into their ppm equivalent (mg/L or mg/kg) (Table S8). In case multiple toxicity concentration values were reported for a given stressor-species pair in ECOTOX, we selected the minimum concentration value for analysis. This process yielded toxicity concentration values for 80 PHs across 221 species (Table S8). Thereafter we constructed a stressor-species network for these 80 PHs, where the edge between a stressor and a species is represented by the logarithm of the standardized toxicity concentration value. Further, we visualized a subnetwork of this stressor-species network for PAHs from the EPA priority PAHs list (EPA, 1979; Keith, 2015) in Cytoscape (Shannon et al., 2003).

Similarly, to construct the stressor-species network using the bioconcentration factor value, we first retrieved the bioconcentration factor values for the curated list of 320 PHs across different species. Note that bioconcentration factor values for stressor-species pairs are provided in the L/kg unit in ECOTOX. In case multiple bioconcentration factor values were reported for a given stressor-species pair in ECOTOX, we selected the maximum bioconcentration factor value for analysis. This process yielded bioconcentration factor values for 28 PHs across 59 species (Table S9) for which we constructed a stressor-species network, where the edge between a stressor and a species is represented by the logarithm of the bioconcentration factor value. Finally, we visualized a subnetwork of this stressor-species network for PAHs from the EPA priority PAHs list (EPA, 1979; Keith, 2015) in Cytoscape (Shannon et al., 2003).

### 2.6. Construction of Species Sensitivity Distributions for petroleum hydrocarbons

Species Sensitivity Distribution (SSD) is a commonly used tool for performing ecotoxicological risk assessment (Kamo, 2023; Kooijman, 1987; van Straalen and Denneman, 1989). SSD utilizes statistical distribution of responses to chemical exposure by various species (Posthuma et al., 2001) and provides a threshold chemical concentration (HC05) which is hazardous to 5% of species but is protective of 95% of species in a particular environment (Dowse et al., 2013; EPA, 1998). In order to construct the SSDs for PHs, we followed the steps outlined in the SSD Toolbox technical manual (Etterson, 2020) to curate data from ECOTOX (Figure 2).

**Figure 2:**
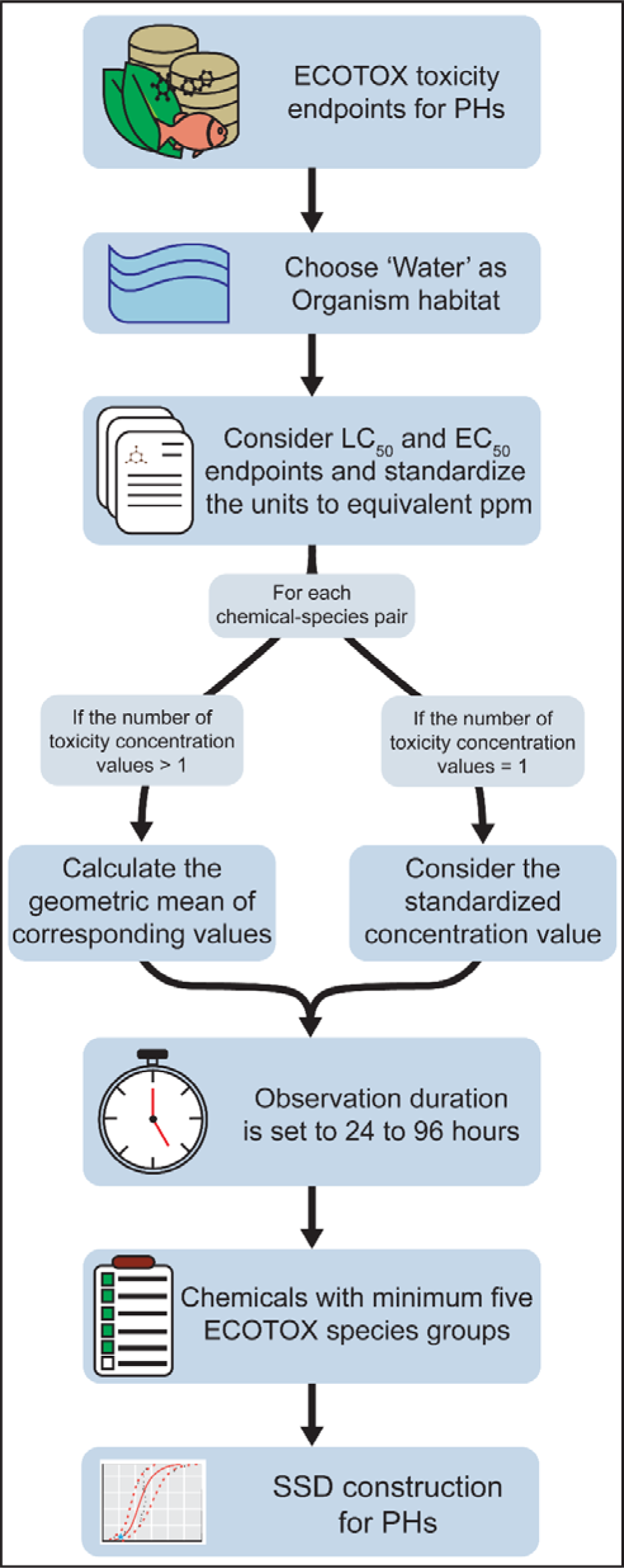
Workflow to curate chemical acute toxicity endpoint data from ECOTOX database to construct species sensitivity distributions for petroleum hydrocarbons.

First, we selected toxicity data only for aquatic species by choosing the organism habitat as ‘Water’ (Figure 2). We then retrieved chemical concentration values for acute toxicity endpoints (LC_50_ and EC_50_) of PHs from ECOTOX along with information on their associated species (Figure 2). Next, we standardized these concentration values by converting their corresponding units to ppm equivalents (mg/L or mg/kg). In the case of multiple toxicity values for a given chemical-species pair, we considered the geometric mean (Schwarz and Tillmanns, 2019; Wheeler et al., 2002) of the standardized concentration values and computed the logarithm of this mean (Figure 2). To ensure the relevance of the data, we excluded any data points with a mean observation time less than 24 hours or exceeding 96 hours (EPA, 2002), retaining only toxicity concentrations from acute toxicity studies. Finally, we considered PHs with toxicity data reported across species belonging to at least five different ECOTOX species groups to construct SSDs (Figure 2) (Belanger et al., 2017).

We relied on two different tools namely, SSD Toolbox developed by the US EPA (https://www.epa.gov/comptox-tools/species-sensitivity-distribution-ssd-toolbox) and R-based ssdtools package developed by the Ministry of Environment and Climate Change Strategy of British Columbia (Schwarz and Tillmanns, 2019) to construct SSDs for PHs. In both cases, we employed the maximum likelihood method to fit five statistical distributions, namely, Log-Normal, Log-Logistic, Log-Gumbel, Weibull and Burr Type III to the toxicity concentration data. The SSD Toolbox and ssdtools utilize bootstrap resampling to quantify uncertainty in fitted parameters and estimate confidence intervals (Etterson, 2020; Schwarz and Tillmanns, 2019). In this study, we set the number of bootstrap resampling iterations to 10,000.

## 3. Results and discussion

### 3.1. Exploration of the curated list of petroleum hydrocarbons

Petroleum hydrocarbons (PHs) are a class of organic compounds that are composed mainly of carbon and hydrogen, and originate from crude oils (Kuppusamy et al., 2020). In this study, we curated a list of 320 petroleum hydrocarbons (PHs) that are experimentally detected in 11 different fuel oils including crude oil (Methods; Figure 1a; Table S1). For these PHs, we first obtained their chemical structure from PubChem, and then employed RDKit (https://www.rdkit.org) to classify them into aliphatic and aromatic compounds. Among the 320 PHs, we identified 117 as aliphatic hydrocarbons (no aromatic ring), 60 as monocyclic aromatic hydrocarbons (one aromatic ring), and 83 as polycyclic aromatic hydrocarbons (more than one aromatic ring) (Table S1). Further, we explored the distribution of these 320 PHs across the 11 fuel oils and observed that more than 100 PHs are found in two fuel oils namely, crude oil and gasoline (Figure 1b).

Next, we checked the presence of these 320 PHs in various chemical regulation lists, namely, the United States high production volume (USHPV) (https://comptox.epa.gov/dashboard/chemical-lists/EPAHPV) chemical list, Organisation for Economic Co-operation and Development high production volume (OECD HPV) (https://hpvchemicals.oecd.org/ui/Search.aspx) chemical list, substances of very high concern (SVHC) (https://echa.europa.eu/candidate-list-table) and REACH prohibited chemicals list (Figure 1c). We observed that 49 PHs are documented to be produced in high volumes globally, 27 PHs are substances of very high concern and 21 PHs are prohibited for use under REACH regulation (Figure 1c). Moreover, we noted that the majority of HPV and SVHC chemicals among the PHs are classified as aliphatic PHs (Figure 1c).

The Chemical and Products Database (CPDat) (Dionisio et al., 2018) is a US EPA project that has compiled information on 75,000 chemicals and their presence in 15,000 consumer products. CPDat provides product use categories (PUCs) and functional use information for these chemicals across products. We leveraged data within CPDat to check the presence of the 320 PHs across various PUCs and their reported functional use information (Figure 1d,e). We retrieved PUC data for 54 PHs across 17 categories, with 20 PHs classified under the categories ‘Vehicle’ and ‘Home maintenance’ (Figure 1d). We also retrieved functional use data for 61 chemicals across 76 different uses, with more than 20 PHs reported to be used as ‘solvent’ and ‘fragrance component’ (Figure 1e).

The US EPA has identified 16 polycyclic aromatic hydrocarbons (PAHs) as priority pollutants due to their frequent occurrence in environmental samples such as air, water, soil, and food, and their potential carcinogenic and mutagenic properties (Eom et al., 2007; EPA, 1979; Sharma et al., 2007; Sopian et al., 2021; Yan et al., 2004; Zelinkova and Wenzl, 2015; Zhang and Tao, 2009; Zhuo et al., 2017). These priority PAHs include naphthalene, acenaphthene, fluorene, phenanthrene, anthracene, fluoranthene, pyrene, benzo[a]anthracene, chrysene, benzo[b]fluoranthene, benzo[k]fluoranthene, benzo[a]pyrene, benzo[ghi]perylene, indeno[1,2,3-cd]pyrene and dibenzo[a,h]anthracene (EPA, 1979). We observed that 15 of these 16 priority PAHs are present in the curated list of 320 PAHs (Table S1). Dibenzo[a,h]anthracene was not included in the curated PH list as it did not have an associated fuel oil source.

### 3.2. Stressor-AOP network for PHs

A stressor-AOP network can elucidate different AOPs associated with a stressor of interest, thereby enhancing our understanding of the various adverse biological effects induced by that stressor (Aguayo-Orozco et al., 2019; Sahoo et al., 2024a). In this study, we leveraged the ecotoxicologically-relevant biological endpoints from three different sources namely, CTD (Davis et al., 2023), ToxCast (Dix et al., 2007) and ECOTOX (Olker et al., 2022), and identified 206 KEs from 177 ecotoxicologically-relevant AOPs to be associated with 75 of the 320 PHs (Methods; Table S5). Thereafter we mapped a PH to an ecotoxicologically-relevant AOP if at least one KE within that AOP is associated with the PH. Following this procedure, we identified 3265 PH-AOP associations for 75 PHs and 177 ecotoxicologically-relevant AOPs, and constructed a bipartite stressor-AOP network, which we designate as ‘PH-AOP’ network (Table S7). Notably, all the PH-AOP associations in the constructed stressor-AOP network are identified through the systematic data integrative approach followed in this study and none were documented in AOP-Wiki.

Next, we computed the coverage score for all PH-AOP links in the constructed stressor-AOP network (Methods) and observed that benzo[a]pyrene (CAS: 50-32-8) is associated with all the KEs (coverage score = 1) in two AOPs namely, AOP: 30 and AOP: 263. Further, we computed the levels of relevance for all the PH-AOP associations and observed that 548 links between 31 PHs and 122 AOPs are classified as Level 1, 2578 links between 75 PHs and 110 AOPs are classified as Level 2, 77 links between 19 PHs and 34 AOPs are classified as Level 3, and 62 links between 10 PHs and 33 AOPs are classified as Level 5 (Table S7). Note, all the Level 4 links between PHs and AOPs also satisfy Level 5 criterion, and therefore we had no PH-AOP link with Level 4 relevance.

Notably, the constructed PH-AOP network provides 975 stressor-AOP links for 14 priority PAHs and 171 ecotoxicologically-relevant AOPs with varying coverage scores and levels of relevance (Table S7). Here we observed that 305 links between 14 PAHs and 92 AOPs are classified as Level 1, 591 links between 14 PAHs and 99 AOPs are classified as Level 2, 41 links between 8 PAHs and 24 AOPs are classified as Level 3, and 38 links between 4 PAHs and 31 AOPs are classified as Level 5 (Table S7). We noted that, benzo[a]pyrene (CAS: 50-32-8) is associated with the maximum number of ecotoxicologically-relevant AOPs (169), with 29 AOPs identified to have Level 5 stressor-AOP links. Figure 3 shows a portion of the PH-AOP network, comprising Level 5 stressor-AOP links for 10 PHs and 33 AOPs, wherein the 4 PAHs are marked in red border.

**Figure 3:**
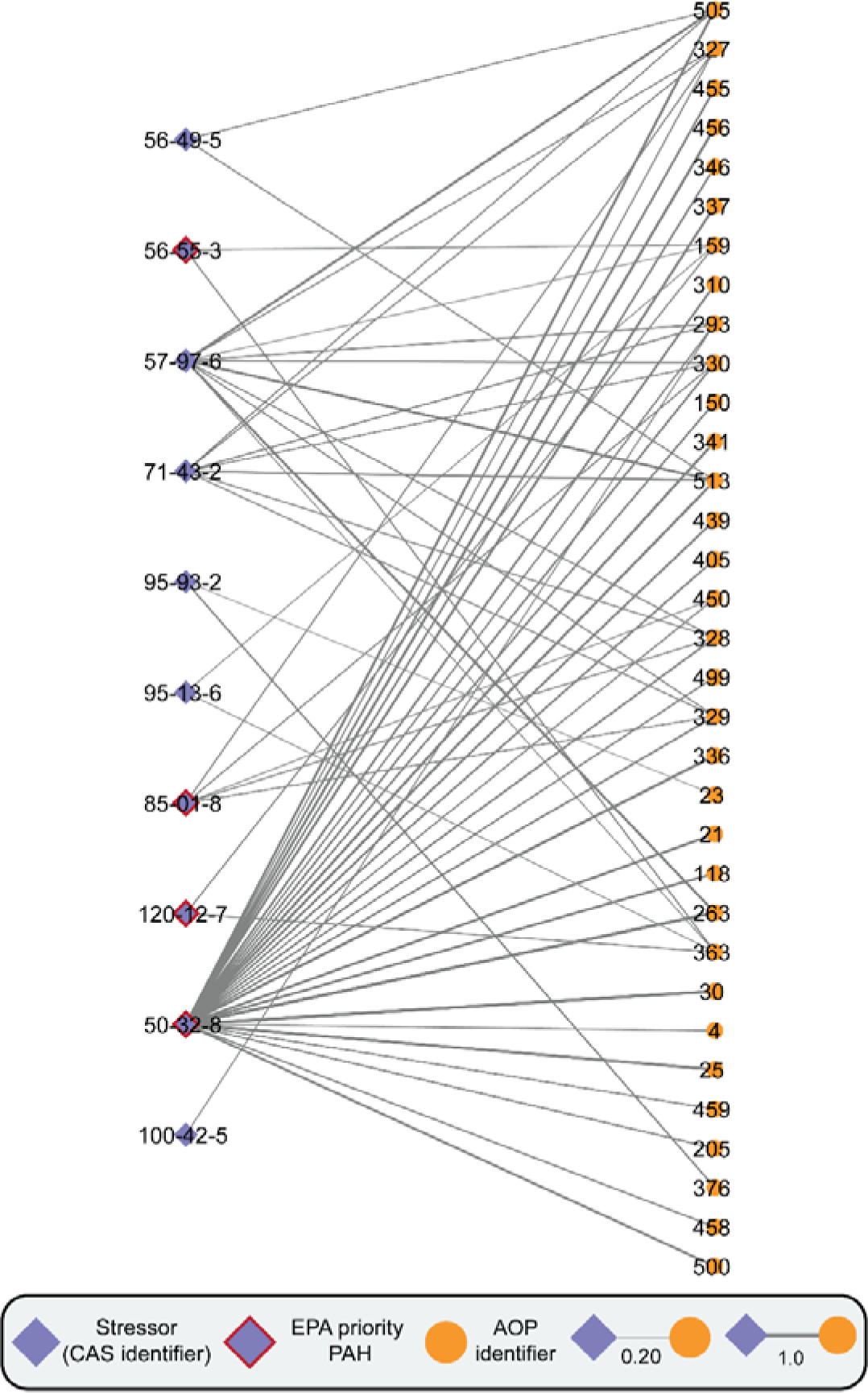
Visualization of a stressor-centric AOP network for PHs. In the stressor-AOP network, only edges or stressor-AOP links with Level 5 relevance are shown. The edges in the stressor-AOP network are weighted based on their coverage score, i.e., the fraction of KEs within AOP that are linked with the PHs. Further, the nodes in the stressor-AOP network that correspond to the EPA priority PAHs are highlighted with red borders.

### 3.3. Exploration of ecotoxicologically-relevant pathways in AOP network associated with Benzo[a]pyrene

PHs are known contaminants in soil and aquatic ecosystems. They can persist in the environment and negatively impact various ecological species (Logeshwaran et al., 2018). Oil spills are frequently documented events that result in the accumulation of these toxic PHs in aquatic environments (Honda and Suzuki, 2020). Benzo[a]pyrene (B[a]P) is a well-documented terrestrial and aquatic pollutant which is found in different fuel oils namely, crude oil, diesel, gasoline, lubricating and motor oils, number 2 fuel oil, and number 6 fuel oil (Table S1). Therefore, we studied the ecotoxicity induced by B[a]P by identifying the highly relevant AOPs associated with B[a]P (designated as B[a]P-AOPs) in the constructed PH-AOP network. We filtered 29 B[a]P-AOPs by selecting stressor-AOP links with Level 5 relevance and coverage score threshold of 0.4 (i.e., ≥ 0.4) in the constructed PH-AOP network.

Further, we computed cumulative weight of evidence (WoE) (Ravichandran et al., 2022; Sahoo et al., 2024b) for each of the 29 B[a]P-AOPs and observed that 11 B[a]P-AOPs have ‘High’ cumulative WoE, 15 B[a]P-AOPs have ‘Moderate’ cumulative WoE (Table S10). Thereafter, we analyzed the ecotoxicity induced by B[a]P by constructing and analyzing an undirected AOP network of the 29 B[a]P-AOPs (Figure S2). We observed that the B[a]P-AOPs form two connected components (wherein at least two B[a]P-AOPs are connected) namely, C1 and C2, and one isolated node, with the largest connected component C1 consisting of 22 B[a]P-AOPs (Figure S2).

Next, we constructed a directed network of the 22 B[a]P-AOPs to explore the interaction among these AOPs (Figure 4). We observed that the directed network comprises 92 KEs and 125 KERs, wherein 51 KEs (including 9 MIEs and 13 AOs) are associated with B[a]P through integration of biological endpoint data from various sources. Thereafter, we analyzed node-centric properties of the directed network by employing various network measures (Table S11). The KE ‘Altered, Cardiovascular development/function’ (KE: 317) and the AO ‘N/A, Breast Cancer’ (KE: 1193) have the highest in-degree of 5. The MIE ‘Activation, AhR’ (KE: 18) has the highest out-degree of 12. The AO ‘Apoptosis’ (KE: 1262) has the highest betweenness centrality value suggesting that this node is centrally located in the network (Figure S3) (Villeneuve et al., 2018). The MIE ‘Activation, AhR’ (KE: 18) has the highest eccentricity value suggesting that this node is the farthest node in the network (Figure S4) (Takes and Kosters, 2011). Further, we utilized Abstract Sifter (Baker et al., 2017) (https://comptox.epa.gov/dashboard/chemical/pubmed-abstract-sifter/) and AOP-helpFinder, an artificial intelligence (AI) based tool (Jaylet et al., 2023; Jornod et al., 2022) (https://aop-helpfinder.u-paris-sciences.fr/), to find associations between B[a]P-induced toxicities and the 92 KEs in the B[a]P-AOP directed network (Table S11).

**Figure 4:**
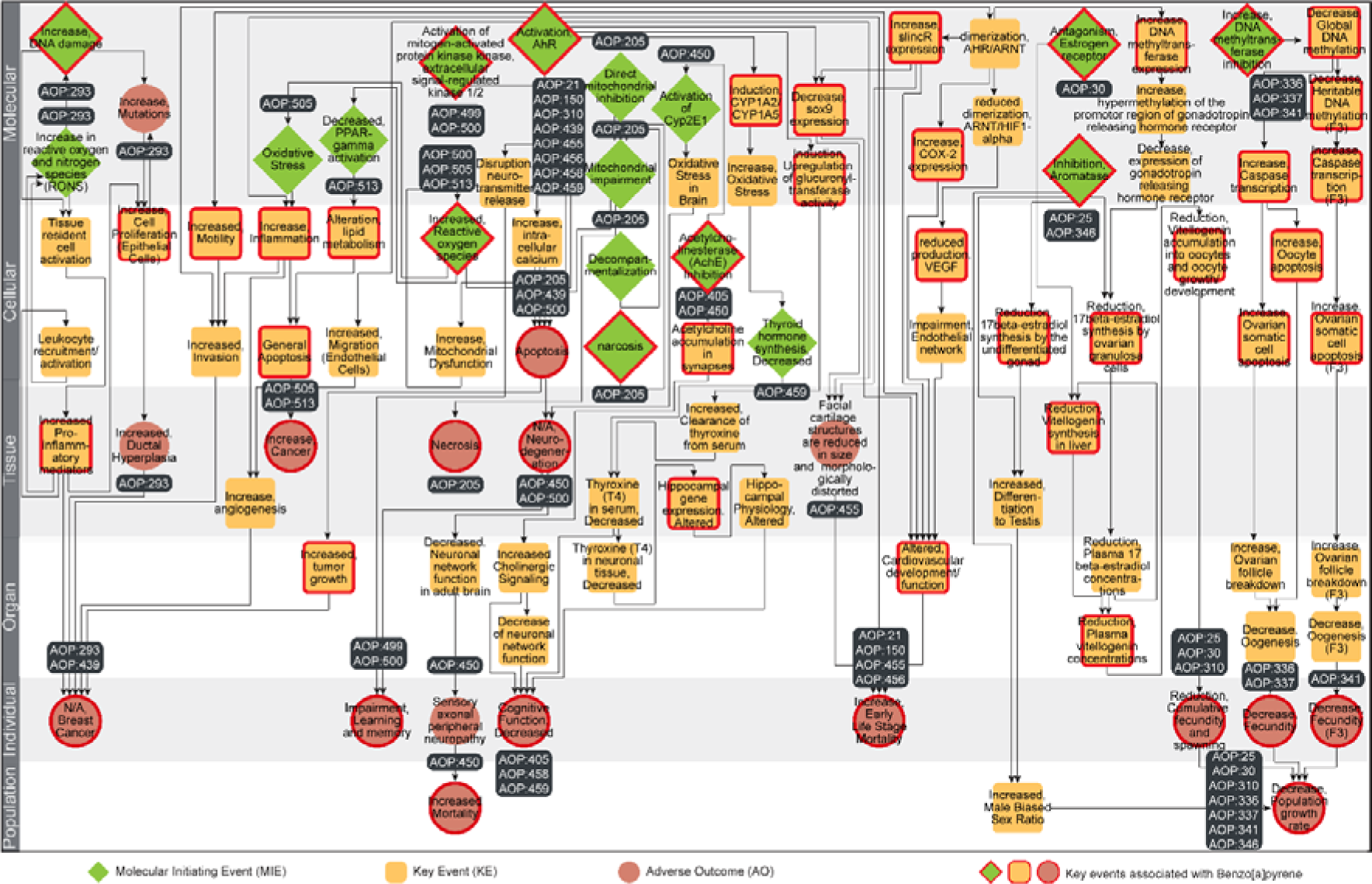
Directed network, corresponding to the largest connected component in the undirected B[a]P-AOP network, comprising 92 KEs and 125 KERs. Among the 92 KEs, 17 are categorized as MIEs (denoted as diamond), 17 are categorized as AOs (denoted as circle), and the remaining 58 are categorized as KEs (denoted as rounded square). The 51 KEs (including MIEs and AOs) associated with B[a]P are marked in ‘red’. In this figure, the 92 KEs are arranged vertically according to their level of biological organization.

#### 3.3.1. Toxicity pathway linking B[a]P exposure to transgenerational effects

Benzo[a]pyrene (B[a]P) is a ubiquitous environmental pollutant that has been associated with transgenerational health consequences in humans and animals (Booc et al., 2014; Pandelides et al., 2023; Wan et al., 2023). Here, we observed a B[a]P-AOP titled ‘DNA methyltransferase inhibition leading to transgenerational effects (2)’ (AOP: 341) having a cumulative WoE of ‘Moderate’ (Table S10) and taxonomical relevance to *Daphnia magna* (Table S2). Therefore, we relied on this ecotoxicologically-relevant AOP to understand the rationale behind B[a]P-induced transgenerational effects.

Different *in vitro* experiments showed a reduction in methyltransferase reactions in embryonic fibroblasts upon exposure to B[a]P (Wojciechowski and Meehan, 1984; Yauk et al., 2008). Subsequently, Corrales *et al*. (Corrales et al., 2014) observed a decrease in global DNA methylation following a parental and continued embryonic waterborne B[a]P exposure in zebrafish embryo and larvae. Wan *et al*. (Wan et al., 2023) showed that ancestral B[a]P exposure in medaka fish led to transgenerational skeletal deformities and changes in gene expression, primarily mediated by histone modifications and miRNAs rather than DNA methylation. Furthermore, Lin *et al*. showed that maternal exposure to B[a]P increased oxidative stress, leading to higher expression of cleaved caspase-3 in the neuroepithelium of mice embryos (Lin et al., 2018). Malott *et al*. observed that the gestational exposure to B[a]P led to ovarian follicle depletion in the mice offspring ovaries and oocytes, with increased mitochondrial superoxide levels and induced apoptosis via the mitochondrial pathway (Malott et al., 2022). Finally, Sui *et al*. observed that B[a]P exposure compromised oogenesis in mice offspring, leading to reduced oocyte maturation, increased meiotic abnormalities, and decreased embryo developmental competence due to mitochondrial dysfunction, oxidative stress, and early apoptosis, all of which led to reduced population sizes in later generations (Sui et al., 2020). Thus, we were able to explore a potential toxicity pathway underlying B[a]P-induced transgenerational effects by leveraging various published evidence.

### 3.4. Stressor-species networks for PHs

#### 3.4.1. Using toxicity concentration as edge weight

A stressor-species network constructed using toxicity concentrations as edge weights can provide information on the variability of chemical toxicity across different species (Wang et al., 2024). In this study, we leveraged acute toxicity concentration data (LC_50_ and EC_50_) for PHs from the ECOTOX database and constructed a bipartite stressor-species network (Methods; Table S8). The resulting network comprises 80 PHs and 221 species (spanning 12 ECOTOX species groups) with 815 stressor-species links (Table S8). We observed that the 32 PAHs in the stressor-species network are documented to be toxic to the highest number of species (163), spanning 11 ECOTOX species groups (Figure 5). The species groups most tested by the PAHs are ‘Crustaceans’, ‘Fish’, and ‘Algae’ (Figure 5). Figure 6 shows the stressor-species network for 14 priority PAHs, which are linked to 160 species through 350 stressor-species connections. Among the 14 PAHs, fluoranthene (CAS: 206-44-0) has been documented to be toxic to the highest number of species (75), followed by naphthalene (CAS: 91-20-3) to 55 species, and phenanthrene (CAS: 85-01-8) to 50 species (Figure 6). The species *Daphnia magna* and *Oncorhynchus mykiss* are linked to 12 and 11 PAHs, respectively (Figure 6), making them the most tested species by the priority PAHs. Overall, we observed that ‘Crustaceans’ are the most tested ECOTOX species group by the priority PAHs.

**Figure 5:**
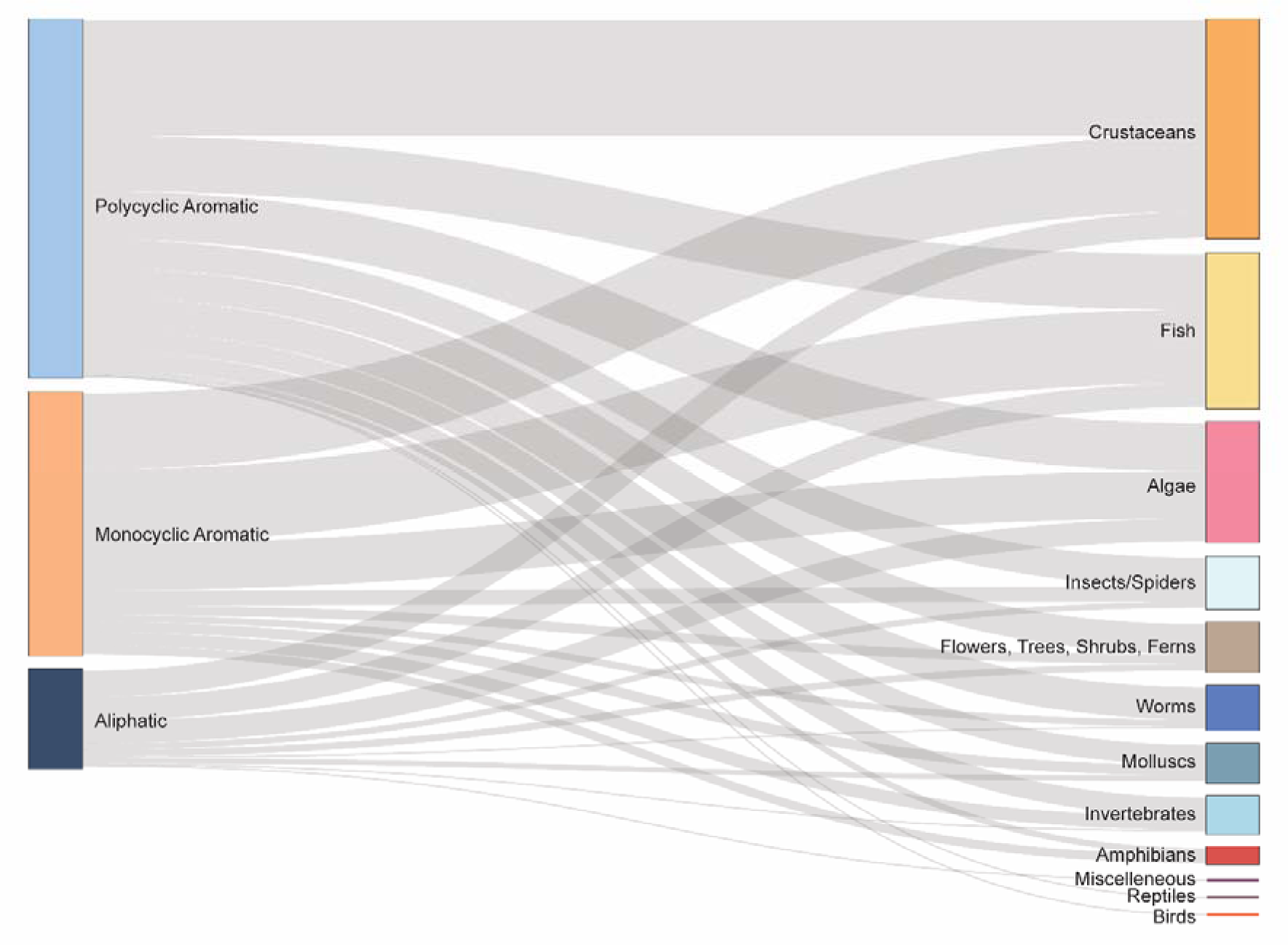
Sankey plot depicting associations between different types of petroleum hydrocarbons and ECOTOX species groups through their acute toxicity concentration data (LC_50_ and EC_50_). The plot provides associations between 3 types of PHs (aliphatic, monocyclic aromatic and polycyclic aromatic) with 12 ECOTOX species groups.

**Figure 6:**
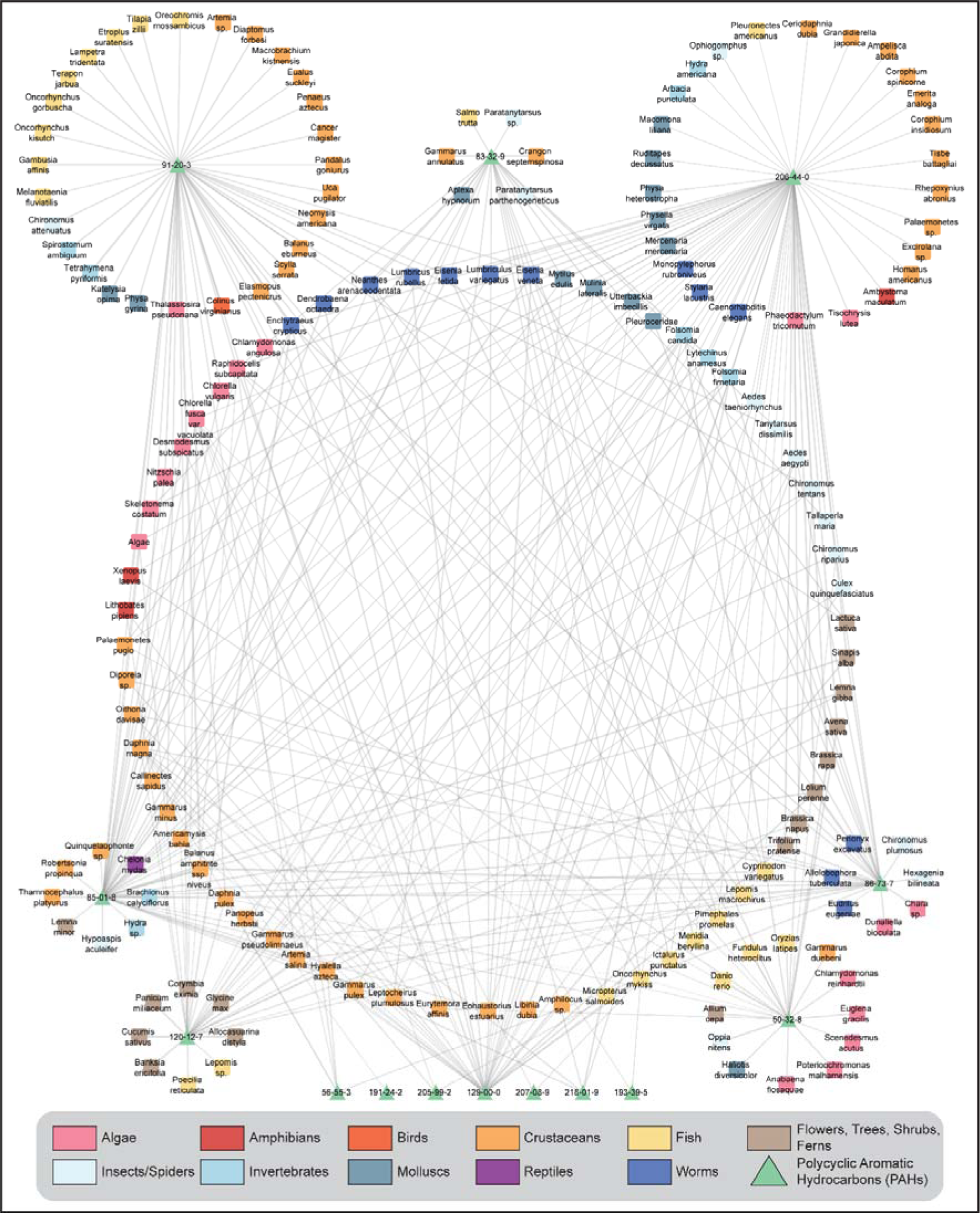
Stressor-species network constructed for EPA priority polycyclic aromatic hydrocarbons (PAHs) using the acute toxicity concentration as edge weights. The network comprises 14 priority PAHs and 160 species with 350 stressor-species links. The edges in the stressor-AOP network are represented by the logarithm of standardized acute toxicity concentration data (LC_50_ and EC_50_). The species in the stressor-species network are classified according to the ECOTOX species groups.

#### 3.4.2. Using bioconcentration factor as edge weight

A stressor-species network constructed using the bioconcentration factor as edge weight can elucidate the extent of chemical absorption by various species from their environment through respiration and dermal surfaces, excluding absorption through diet (Arnot and Gobas, 2006). In this study, we leveraged the bioconcentration factors for PHs from the ECOTOX database and constructed a bipartite stressor-species network (Methods; Table S9). The resulting network comprises 28 PHs and 59 species (spanning 9 ECOTOX species groups) with 159 stressor-species links (Table S9). We observed that 22 PAHs in the stressor-species network are documented to be absorbed by majority of the ECOTOX species groups (Figure 7). Species groups namely, ‘Crustaceans’, ‘Fish’ and ‘Molluscs’ are reported to absorb 21, 14, and 12 PHs, respectively (Table S9). Figure 8 shows the stressor-species network of 13 priority PAHs with 54 species comprising 117 stressor-species links. Among the 13 PAHs, benzo[a]pyrene (CAS: 50-32-8) is documented to be absorbed in highest number of species (20), followed by phenanthrene (CAS: 85-01-8) and fluoranthene (CAS: 206-44-0) in 17 species each. We observed that many of the PAHs are documented to be absorbed in species namely, *Daphnia magna*, *Daphnia pulex* and *Hyalella azteca* from ‘Crustaceans’ ECOTOX species group (Figure 8).

**Figure 7:**
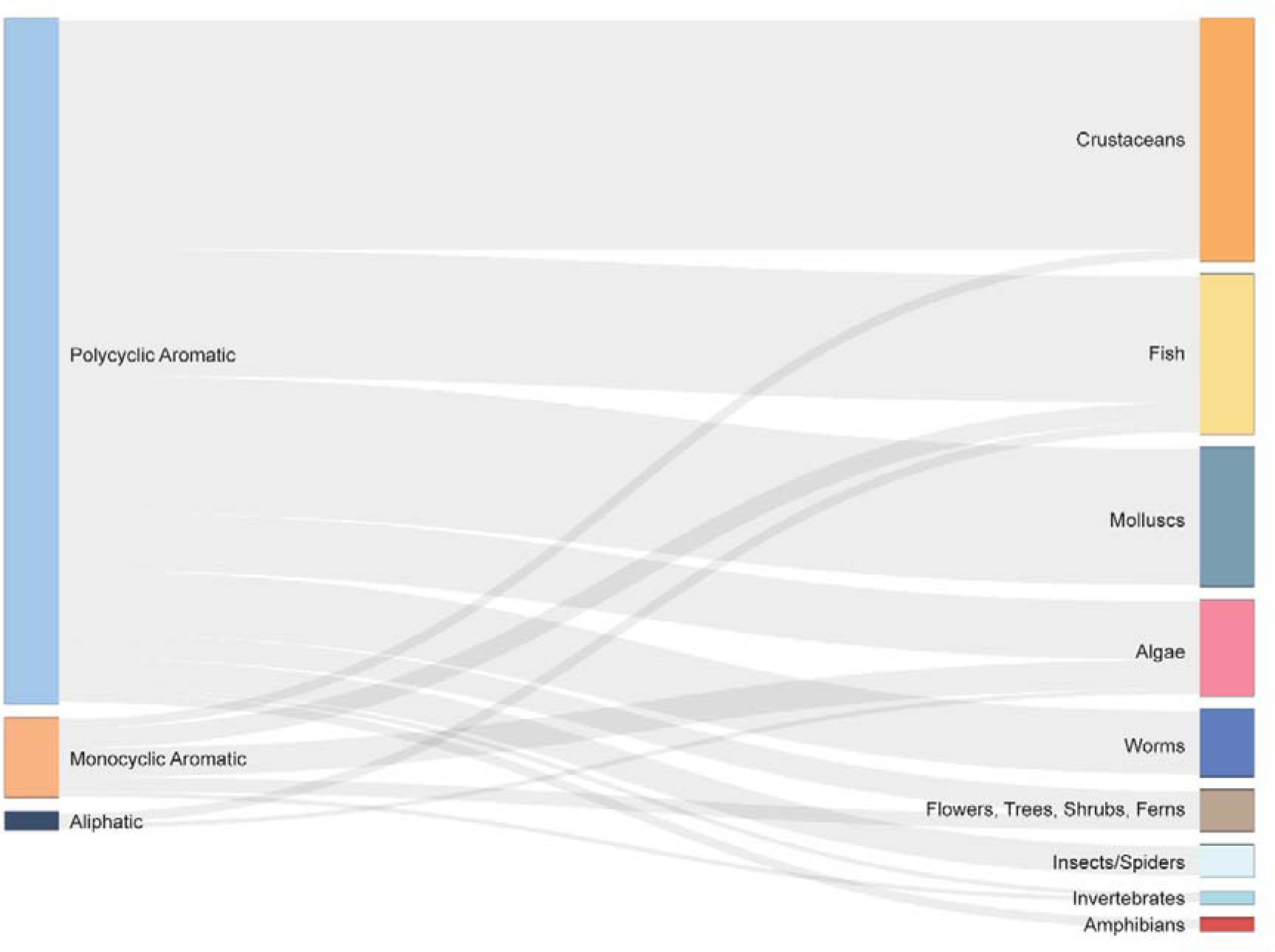
Sankey plot depicting associations between different types of petroleum hydrocarbons and ECOTOX species groups through their bioconcentration factors. The plot provides associations between 3 types of PHs (aliphatic, monocyclic aromatic and polycyclic aromatic) with 9 ECOTOX species groups.

**Figure 8:**
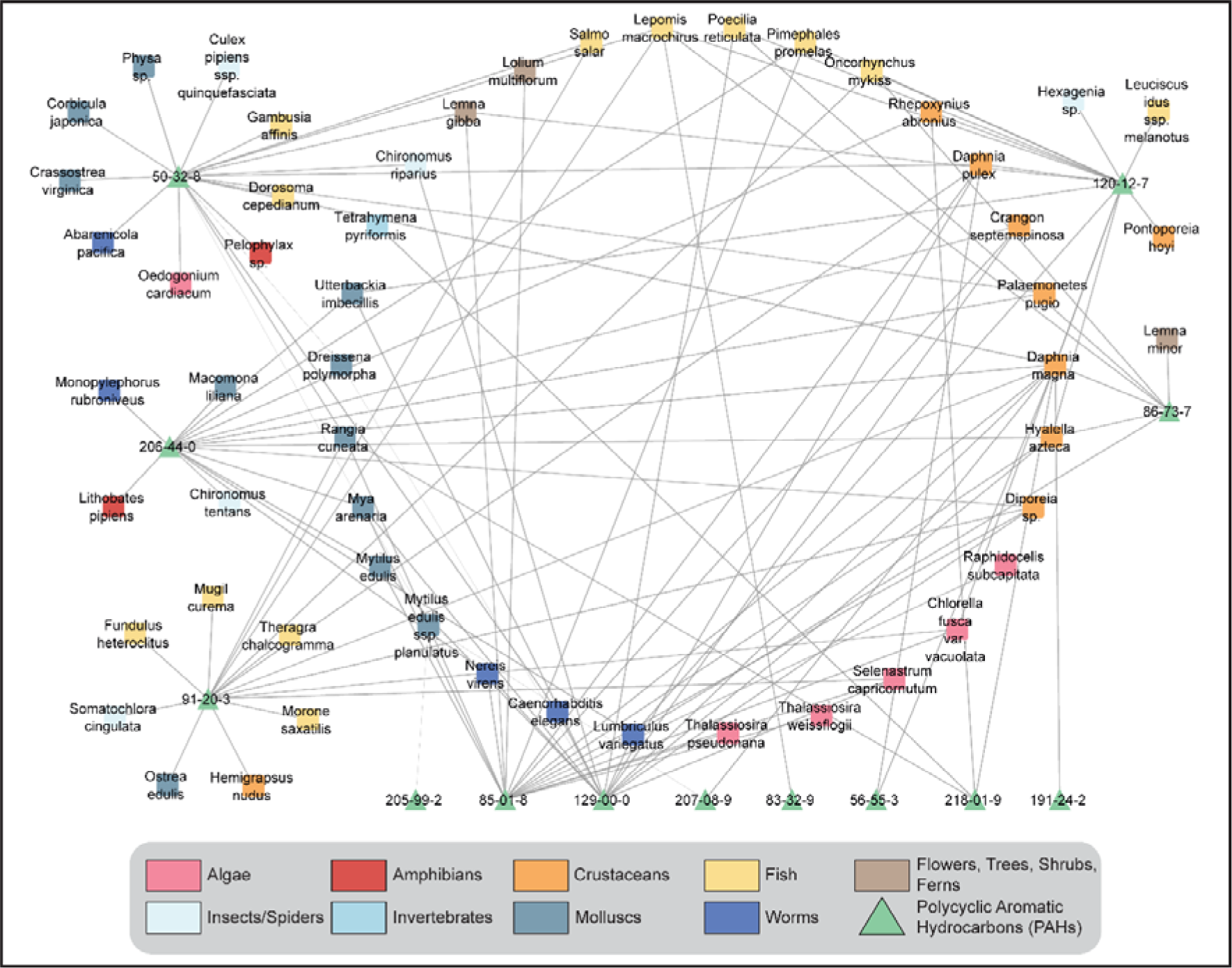
Stressor-species network constructed for EPA priority polycyclic aromatic hydrocarbons (PAHs) using the bioconcentration factors as edge weights. The network comprises 13 priority PAHs, 54 species and 117 stressor-species links. The edges in the stressor-AOP network are represented by the logarithm of bioconcentration factors. The species in the stressor-species network are classified according to the ECOTOX species groups.

### 3.5. Species sensitivity distribution for EPA priority PAHs

Species sensitivity distribution (SSD) has been extensively used to derive environmental quality criterion or hazard concentration of chemicals in different environments (Karthikeyan et al., 2021; Méndez-Fernández et al., 2019; Posthuma et al., 2001). In this study, we relied on the acute toxicity endpoints provided by ECOTOX to construct the SSDs for the priority PAHs (EPA, 1979; Keith, 2015), and leveraged them to derive the corresponding hazard concentrations in aquatic environments (Methods; Figure 2; Table S12). We observed that the acute toxicity endpoints associated with 8 of the 16 priority PAHs namely, fluoranthene, naphthalene, phenanthrene, benzo[a]pyrene, acenaphthene, pyrene, anthracene, and 9H-fluorene are reported in terms of LC_50_ or EC_50_ values, and observed within the duration of 24 to 96 hours in at least 5 ECOTOX species groups (Methods; Figure 2; Table S12). Therefore, we accessed the acute toxicity endpoints from ECOTOX for each of these eight priority PAHs and employed both US EPA SSD Toolbox (Etterson, 2020) and the R-based ssdtools (Schwarz and Tillmanns, 2019) to construct SSDs and derive the corresponding hazard concentration values (Methods).

The SSD Toolbox and ssdtools fit the data using five distributions, namely Log-Normal, Log-Logistic, Log-Gumbel, Weibull and Burr Type III, and provide the HC05 value based on the best-fit model determined by the minimum corrected Akaike Information Criterion (AICc) value (Cavanaugh and Neath, 2019; Etterson, 2020; Schwarz and Tillmanns, 2019). We observed that both tools identified the same best-fit model for each of the eight priority PAHs, and the derived HC05 values were also similar (Figure S5; Table S13). For example, Figure 9 shows the plots of SSD for B[a]P computed using the best-fit Weibull model, as determined by both the SSD Toolbox and ssdtools. Figures S6-S12 present the plots of SSD computed using the best-fit models for each of the other seven priority PAHs, as determined by both the SSD Toolbox and ssdtools.

**Figure 9:**
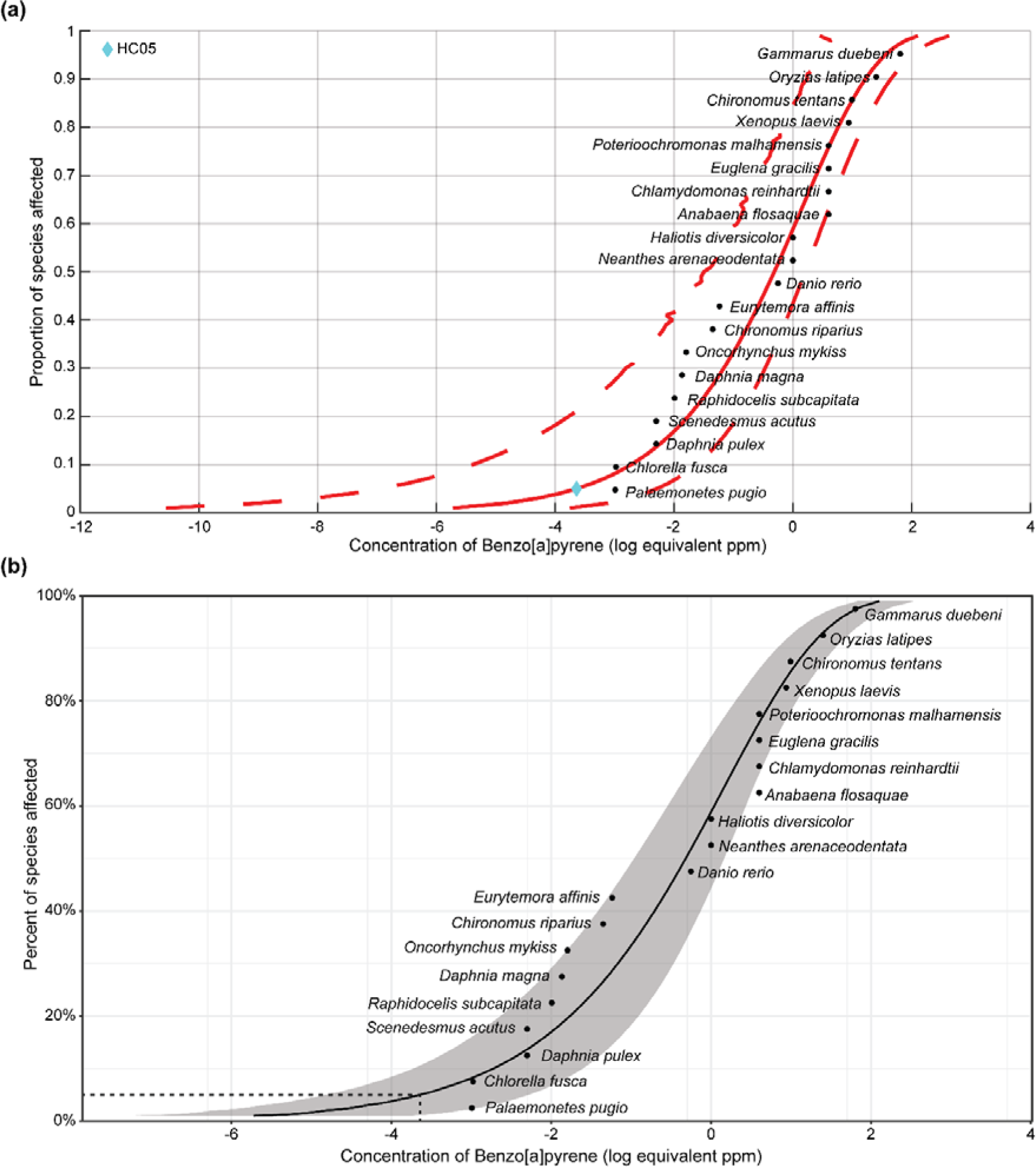
The plots of SSD for benzo[a]pyrene computed using the best-fit Weibull model. **(a)** As determined by US EPA SSD Toolbox where the HC05 value is denoted by cyan colored diamond. **(b)** As determined by ssdtools where the HC05 value is denoted by a dotted line.

Recently, the method of model averaging has been proposed to compute SSDs, wherein the averaged model is obtained by assigning model weight to each of the individual distributions (Fox et al., 2021; Schwarz and Tillmanns, 2019). The HC05 values computed based on model averaging have been observed to be more reliable and stable compared to the values obtained from individual models (Fox et al., 2021; Schwarz and Tillmanns, 2019). Therefore, we computed model-averaged HC05 values for each of the eight priority PAHs and observed that both tools provided similar results (Table 1). We observed that B[a]P has the lowest model-averaged HC05 value, and naphthalene has the highest model-averaged HC05 value (Table 1).

**Table 1:** Model averaged HC05 values and the corresponding standard errors computed by both US EPA SSD Toolbox and ssdtools for eight priority PAHs. The values are given in equivalent ppm units.

| Serial Number | Chemical name | US EPA SSD Toolbox |  | ssdtools |  |
| --- | --- | --- | --- | --- | --- |
|  |  | Model averaged HC05 value | Model averaged HC05 standard error | Model averaged HC05 value | Model averaged HC05 standard error |
| 1 | Fluoranthene | 0.010708 | 0.002079 | 0.010712 | 0.002074 |
| 2 | Naphthalene | 0.69307 | 0.20508 | 0.693079 | 0.19385 |
| 3 | Phenanthrene | 0.053411 | 0.024022 | 0.053633 | 0.02168 |
| 4 | Benzo[a]pyrene | 0.000659 | 0.003752 | 0.000659 | 0.003746 |
| 5 | Acenaphthene | 0.22909 | 0.11273 | 0.22909 | 0.102988 |
| 6 | Pyrene | 0.003423 | 0.002693 | 0.003424 | 0.002594 |
| 7 | Anthracene | 0.00365 | 0.003635 | 0.003678 | 0.003424 |
| 8 | 9H-fluorene | 0.25044 | 0.17644 | 0.252645 | 0.166426 |

The HC05 values indicate the toxic effects of chemicals across species, with smaller HC05 values implying greater toxicity (Yanagihara et al., 2022). Based on the model-averaged HC05 values, we ordered the eight priority PAHs from most toxic to least toxic for aquatic organisms as follows: benzo[a]pyrene > pyrene > anthracene > fluoranthene > phenanthrene > acenaphthene > 9H-fluorene > naphthalene. Previously, Chen *et al*. leveraged chronic aquatic toxicity data and observed a similar order of the eight PAHs based on their computed HC05 values (Chen et al., 2020). Moreover, we observed that this decreasing order coincides with the number of benzene rings in these compounds, potentially indicating a correlation between the number of rings in PAHs and their level of toxicity (Wang et al., 2024).

The species located in the region of lower toxicity in the SSD are identified as sensitive to that particular chemical (Karthikeyan et al., 2021). We observed that the species belonging to the ECOTOX species groups ‘Crustaceans’, ‘Fish’ and ‘Molluscs’, are commonly found in the region of lower toxicity in the SSDs computed for the eight PAHs. In particular, the species *Palaemonetes pugio* (crustacean)*, Oncorhynchus mykiss* (fish)*, Americamysis bahia* (crustacean)*, Daphnia pulex* (crustacean), and *Utterbackia imbecillis* (mollusc) were sensitive to more than one PAH. Notably, these species are also connected with the PAHs in the stressor-species network constructed based on bioconcentration factors (Figure 8). Furthermore, we observed *P. pugio* is highly sensitive to benzo[a]pyrene and fluoranthene, and *A. bahia* is highly sensitive to phenanthrene and pyrene.

In a nutshell, we leveraged acute toxicity endpoints from ECOTOX to construct SSDs for the eight priority PAHs and derived the corresponding HC05 values using the model averaging method in both US EPA SSD Toolbox and ssdtools. The derived HC05 values were similar across both tools and helped identify a toxicity order for the eight PAHs.

## 4. Conclusion

Petroleum hydrocarbons are released into the environment through various human activities or accidental oil spills, where they can persist and pose long-term ecological risks. Thus, it is imperative to study the effects of PH exposure on species inhabiting the contaminated environments. In this study, we relied on the reports published by the Total Petroleum Hydrocarbon Criteria Working Group (TPHCWG) (Gustafson et al., 1997; Potter and Simmons, 1998) and curated a list of 320 PHs that were experimentally identified to be present in various fuel oils. We utilized this list to explore the mechanism of PH-induced ecotoxicity, and specifically focusing on the 16 EPA priority PAHs, assessed their risks in aquatic environments. First, we curated a list of 195 ecotoxicologically-relevant AOPs from AOP-Wiki. Subsequently, we systematically integrated biological endpoint data from three sources namely, ToxCast, CTD and ECOTOX, and identified 206 KEs from 177 ecotoxicologically-relevant AOPs to be associated with 75 of 320 PHs. We constructed a stressor-centric AOP network comprising these 75 PHs and 177 ecotoxicologically-relevant AOPs linked through 3265 edges with varying levels of relevance and coverage scores. We leveraged this stressor-AOP network and identified 29 AOPs relevant for benzo[a]pyrene-induced toxicity, constructed an AOP network, and performed a case-study to understand its transgenerational effects in ecological species. Next, we utilized the acute toxicity data within ECOTOX, constructed a stressor-species network comprising 80 PHs linked to 221 species, and observed that ‘Crustaceans’ species group was documented to be affected by many of these PHs. Similarly, we utilized the bioconcentration factors data within ECOTOX, constructed a stressor-species network comprising 28 PHs linked to 59 species, and observed that ‘Crustaceans’ species group was documented to bioaccumulate many of these PHs. Finally, we utilized the acute toxicity data within ECOTOX available for eight EPA priority PAHs, constructed their species sensitivity distribution (SSD), and derived corresponding hazard concentrations (HC05) that protect 95% of the species in aquatic environments. In sum, the present study utilizes various network-based approaches along with ecotoxicological data to elucidate PH-induced toxicities and the associated risks for ecological species.

However, the scope of the present study is restricted by several limitations on the available data. The curated list of PHs may be incomplete due to the limitations of current analytical methods in determining the full molecular composition of fuel oils (Bierkens and Geerts, 2014). We observed that many of the AOPs within AOP-Wiki have no taxonomic applicability annotation thereby leading to the curation of an incomplete set of ecotoxicologically-relevant AOPs. Further, we observed that the derived HC05 value for some of the EPA priority PAHs have wider confidence intervals implying the need of more stressor-specific toxicity studies across diverse ecological species in order to accurately derive the threshold concentrations (Machin et al., 2000). Importantly, the derived threshold concentrations of the priority PAHs may not be applicable to a specific aquatic environment as the information on the test location for the underlying toxicity data was sparse.

Nonetheless, this study advances our understanding of the ecotoxicological effects of individual PHs through network-based approaches. The stressor-AOP network constructed using ecotoxicologically-relevant endpoints for PHs has facilitated the investigation of toxicity pathways leading to PH-specific adverse outcomes. Further, the inferences from this study using acute toxicity data can aid in assessing the risks associated with events such as oil spills, which result in a sudden increase in PH concentrations in ecosystems. In future, the derived threshold concentrations of chemicals in conjunction with concentrations necessary to trigger the KEs in the associated AOP can be employed to assess environmental quality standards. In conclusion, this study explores the ecotoxicological effects and risks associated with PH exposure in ecosystems, thereby assisting in their regulation and enabling the formulation of effective mitigation and remediation strategies for various PH contaminations.

## Data availability

The data associated with this study is contained in the article or in the supplementary information files.

## Acknowledgements

Areejit Samal would like to acknowledge funding from the Department of Atomic Energy (DAE), Government of India via Apex project to The Institute of Mathematical Sciences (IMSc) Chennai. The funders have no role in study design, data collection, data analysis, manuscript preparation or decision to publish.

## Author contributions

**Conceptualization:** Ajaya Kumar Sahoo, Shreyes Rajan Madgaonkar, Nikhil Chivukula, Shambanagouda Rudragouda Marigoudar, Krishna Venkatarama Sharma, Areejit Samal; **Methodology:** Ajaya Kumar Sahoo, Shreyes Rajan Madgaonkar, Nikhil Chivukula, Panneerselvam Karthikeyan, Shambanagouda Rudragouda Marigoudar, Krishna Venkatarama Sharma, Areejit Samal; **Data curation:** Ajaya Kumar Sahoo, Shreyes Rajan Madgaonkar, Nikhil Chivukula, Kundhanathan Ramesh; **Formal analysis:** Ajaya Kumar Sahoo, Shreyes Rajan Madgaonkar, Nikhil Chivukula, Panneerselvam Karthikeyan, Kundhanathan Ramesh, Shambanagouda Rudragouda Marigoudar, Krishna Venkatarama Sharma, Areejit Samal; **Visualization:** Ajaya Kumar Sahoo, Shreyes Rajan Madgaonkar, Nikhil Chivukula, Kundhanathan Ramesh; **Software:** Ajaya Kumar Sahoo, Shreyes Rajan Madgaonkar, Nikhil Chivukula; **Writing – original draft:** Ajaya Kumar Sahoo, Shreyes Rajan Madgaonkar, Nikhil Chivukula, Panneerselvam Karthikeyan, Shambanagouda Rudragouda Marigoudar, Krishna Venkatarama Sharma, Areejit Samal; **Writing – review & editing:** Ajaya Kumar Sahoo, Shreyes Rajan Madgaonkar, Nikhil Chivukula, Panneerselvam Karthikeyan, Kundhanathan Ramesh, Shambanagouda Rudragouda Marigoudar, Krishna Venkatarama Sharma, Areejit Samal; **Supervision:** Areejit Samal.

## Supplementary Tables

**Table S1:** This table contains the list of 320 petroleum hydrocarbons (PHs) curated from Total Petroleum Hydrocarbon Criteria Working Group (TPHCWG) volume 2 titled ‘Composition of Petroleum Mixtures’ and volume 3 titled ‘Selection of Representative TPH Fractions Based on Fate and Transport Considerations’. For each PH, the table provides the CAS chemical identifier, PubChem chemical identifier, chemical name, fuel oil(s) in which the PH is reported and the corresponding reference(s). Further, for each PH, the table provides its chemical structure in InChIKey and SMILES format, its molecular weight in g/mol, hydrocarbon type computed in RDKit, chemical Kingdom, chemical Superclass, chemical Class predicted by ClassyFire, its product use category and functional use reported in CPDat, its presence in EPA priority polycyclic aromatic hydrocarbons (PAH) list, its presence as high production volume (HPV) chemical in US HPV or OECDHPV list, its presence in substances of very high concern (SVHC) list, and its presence in REACH list of prohibited chemicals.

**Table S2:** This table contains the curated list of 195 ecotoxicologically-relevant high confidence adverse outcome pathways from AOP-Wiki. For each ecotoxicologically-relevant AOP, the table provides the corresponding information on AOP identifier, AOP title, Organisation for Economic Co-operation and Development (OECD) status, Society for the Advancement of Adverse Outcome Pathways (SAAOP) status, and taxonomic applicability of the AOP.

**Table S3:** This table contains information on the 727 Key Events (KEs) present in the curated list of 195 ecotoxicologically-relevant AOPs. For each KE, the table provides the corresponding information on KE identifier, KE title, level of biological organization, and associated AOP identifier(s) (separated by ‘|’ symbol).

**Table S4:** This table contains information on Key Event Relationships (KERs) present in each of the curated list of 195 ecotoxicologically-relevant AOPs. For each AOP, the table provides the AOP identifier, corresponding KER identifier, upstream KE identifier, downstream KE identifier, MIE(s) among upstream and downstream KEs (separated by ‘|’ symbol), AO(s) among upstream and downstream KEs (separated by ‘|’ symbol), adjacency of KER, weight of evidence of KER, and quantitative understanding of KER. Note that the KER identifiers starting with 10000 were manually assigned by the authors as the KER was mentioned in the AOP page but was not assigned an identifier in AOP-Wiki.

**Table S5:** This table contains information on 206 ecotoxicologically-relevant Key Events (KEs) from 195 ecotoxicologically-relevant AOPs that are associated with petroleum hydrocarbons (PHs). For each KE, the table provides corresponding information on KE identifier, CAS chemical identifier (separated by ‘|’) for associated PH(s), sources from which the associations are inferred (separated by ‘|’ symbol), CTD phenotypes (separated by ‘|’ symbol), CTD diseases (separated by ‘|’ symbol), ToxCast assay endpoints (separated by ‘|’ symbol), ECOTOX measurements and trends [mentioned as measurement:trend] (separated by ‘|’ symbol).

**Table S6:** This table contains information on 8454 chemical-gene-phenotype-disease (CGPD) tetramers constructed for petroleum hydrocarbons (PHs) from Comparative Toxicogenomics Database (CTD). For each tetramer, the table provides the corresponding information on the CAS chemical identifier, NCBI gene identifier, NCBI gene name, phenotype identifier, phenotype name, MESH disease identifier, MESH disease name, and species associated with the chemical-gene link and chemical-phenotype link.

**Table S7:** This table provides edge list for the complete stressor-AOP network for petroleum hydrocarbons (PHs). For each edge in the network, the table provides the CAS identifier for the stressor (PHs), AOP identifier, computed coverage score and level of relevance.

**Table S8:** This table provides edge list for the stressor-species network for petroleum hydrocarbons (PHs) constructed using concentration value for acute toxicity endpoints (LC_50_ and EC_50_). For each edge in the network, the table provides the CAS chemical identifier, chemical name, hydrocarbon type for the PH, Latin name and ECOTOX species group for the species, considered toxicity endpoint, toxicity concentration value, its unit in ECOTOX, factor to convert concentration unit into its ppm equivalent (MW corresponds to the molecular weight of the chemical), concentration value in ppm equivalent unit, and logarithm of concentration value in ppm equivalent unit (rounded up to 6 decimals).

**Table S9:** This table provides edge list for the stressor-species network for petroleum hydrocarbons (PHs) constructed using bioconcentration factors. For each edge in the network, the table provides the CAS chemical identifier, chemical name, hydrocarbon type for the PH, Latin name and ECOTOX species group for the species, bioconcentration factor in L/kg, and logarithm of bioconcentration factor value (rounded up to 6 decimals).

**Table S10:** This table contains information on the evidence and the connected component for each of the 29 benzo[a]pyrene (B[a]P)-AOPs. For each AOP, the table provides the corresponding AOP identifier, computed fraction of KERs with ‘High’ evidence (i.e., F(High)), computed fraction of KERs with ‘Moderate’ evidence (i.e., F(Moderate)), computed fraction of KERs with ‘Low’ evidence (i.e., F(Low)), computed fraction of KERs with ‘Not Specified’ evidence (i.e., F(Not Specified)), computed cumulative weight of evidence (WoE), and the connected component identifier in the undirected AOP network associated with B[a]P.

**Table S11:** This table contains information on the computed network statistics and literature evidence of the association with B[a]P for each of the 92 KEs present in the largest connected component (C1) of the B[a]P-AOP network. For each KE, the table provides the corresponding KE identifier, KE title, level of biological organization, KE type, in-degree value, out-degree value, betweenness centrality value, eccentricity value, convergence information, organism in which B[a]P exposure is studied, study type, dosage information, abbreviated description of association with B[a]P, and the corresponding reference.

**Table S12:** This table contains acute toxicity concentration data used to construct species sensitivity distributions (SSDs) for eight priority PAHs. Further, the table provides information on the CAS chemical identifier, chemical name, Latin name and ECOTOX species group for the species, unit and value of the toxicity concentration of the chemical.

**Table S13:** This table contains information on the derived HC05 values of the eight priority PAHs. For each of these PAHs, the table provides the CAS chemical identifier, chemical name, number of associated species in the constructed SSD, number of ECOTOX species groups in the constructed SSD, HC05 value, HC05 lower confidence limit (LCL), HC05 upper confidence limit (UCL) derived from US EPA SSD Toolbox and ssdtools package for the corresponding best-fit model. The values are given in equivalent ppm units and rounded up to 6 decimal places.

## Supplementary Figures

**Figure S1:** Workflow to filter ecotoxicologically-relevant high confidence adverse outcome pathways (AOPs) from AOP-Wiki by employing computation and manual curation in conjunction. (Adapted from Sahoo *et al*., 2024a).

**Figure S2:** Undirected network of benzo[a]pyrene (B[a]P)-AOPs. Each node corresponds to a B[a]P-AOP and an edge between two nodes denotes that the two AOPs share at least one KE. This undirected network has 2 connected components (wherein at least two nodes are connected) which are labeled as C1, C2, and one isolated node.

**Figure S3:** Directed network corresponding to the largest connected component (C1) in the B[a]P-AOP network, where the KEs (including MIEs and AOs) are colored based on their betweenness centrality values. The 51 KEs (including MIEs and AOs) associated with B[a]P are marked in ‘red’. In this figure, the 92 KEs are arranged vertically according to their level of biological organization.

**Figure S4:** Directed network corresponding to the largest connected component (C1) in the B[a]P-AOP network, where the KEs (including MIEs and AOs) are colored based on their eccentricity values. The 51 KEs (including MIEs and AOs) associated with B[a]P are marked in ‘red’. In this figure, the 92 KEs are arranged vertically according to their level of biological organization.

**Figure S5:** Comparison of the derived HC05 values for each of the eight priority PAHs using the corresponding best-fit model in US EPA SSD Toolbox and ssdtools.

**Figure S6:** The plots of SSD for fluoranthene computed using the best-fit Log-Gumbel model. **(a)** As determined by US EPA SSD Toolbox where the HC05 value is denoted by cyan colored diamond. **(b)** As determined by ssdtools where the HC05 value is denoted by a dotted line.

**Figure S7:** The plots of SSD for naphthalene computed using the best-fit Log-Gumbel model. **(a)** As determined by US EPA SSD Toolbox where the HC05 value is denoted by cyan colored diamond. **(b)** As determined by ssdtools where the HC05 value is denoted by a dotted line.

**Figure S8:** The plots of SSD for phenanthrene computed using the best-fit Log-Gumbel model. **(a)** As determined by US EPA SSD Toolbox where the HC05 value is denoted by cyan colored diamond. **(b)** As determined by ssdtools where the HC05 value is denoted by a dotted line.

**Figure S9:** The plots of SSD for acenaphthene computed using the best-fit Weibull model. **(a)** As determined by US EPA SSD Toolbox where the HC05 value is denoted by cyan colored diamond. **(b)** As determined by ssdtools where the HC05 value is denoted by a dotted line.

**Figure S10:** The plots of SSD for pyrene computed using the best-fit Log-Gumbel model. **(a)** As determined by US EPA SSD Toolbox where the HC05 value is denoted by cyan colored diamond. **(b)** As determined by ssdtools where the HC05 value is denoted by a dotted line.

**Figure S11:** The plots of SSD for anthracene computed using the best-fit Log-Gumbel model. **(a)** As determined by US EPA SSD Toolbox where the HC05 value is denoted by cyan colored diamond. **(b)** As determined by ssdtools where the HC05 value is denoted by a dotted line.

**Figure S12:** The plots of SSD for 9H-fluorene computed using the best-fit Log-Gumbel model. **(a)** As determined by US EPA SSD Toolbox where the HC05 value is denoted by cyan colored diamond. **(b)** As determined by ssdtools where the HC05 value is denoted by a dotted line.

## References

Aguayo-Orozco, A., Audouze, K., Siggaard, T., Barouki, R., Brunak, S., Taboureau, O., 2019. sAOP: linking chemical stressors to adverse outcomes pathway networks. Bioinformatics 35, 5391–5392. 10.1093/bioinformatics/btz570

Alford, J.B., Peterson, M.S., Green, C.C. (Eds.), 2014. Impacts of Oil Spill Disasters on Marine Habitats and Fisheries in North America, 1st ed. CRC Press, Boca Raton. 10.1201/b17633

Al-Hawash, A.B., Dragh, M.A., Li, S., Alhujaily, A., Abbood, H.A., Zhang, X., Ma, F., 2018. Principles of microbial degradation of petroleum hydrocarbons in the environment. Egyptian Journal of Aquatic Research 44, 71–76. 10.1016/j.ejar.2018.06.001

Almeda, R., Wambaugh, Z., Chai, C., Wang, Z., Liu, Z., Buskey, E.J., 2013a. Effects of Crude Oil Exposure on Bioaccumulation of Polycyclic Aromatic Hydrocarbons and Survival of Adult and Larval Stages of Gelatinous Zooplankton. PLoS ONE 8, e74476. 10.1371/journal.pone.0074476

Almeda, R., Wambaugh, Z., Wang, Z., Hyatt, C., Liu, Z., Buskey, E.J., 2013b. Interactions between Zooplankton and Crude Oil: Toxic Effects and Bioaccumulation of Polycyclic Aromatic Hydrocarbons. PLoS ONE 8, e67212. 10.1371/journal.pone.0067212

Ankley, G.T., Bennett, R.S., Erickson, R.J., Hoff, D.J., Hornung, M.W., Johnson, R.D., Mount, D.R., Nichols, J.W., Russom, C.L., Schmieder, P.K., Serrrano, J.A., Tietge, J.E., Villeneuve, D.L., 2010. Adverse outcome pathways: A conceptual framework to support ecotoxicology research and risk assessment. Environmental Toxicology and Chemistry 29, 730–741. 10.1002/etc.34

Arnot, J.A., Gobas, F.A., 2006. A review of bioconcentration factor (BCF) and bioaccumulation factor (BAF) assessments for organic chemicals in aquatic organisms. Environmental Reviews 14, 257–297. 10.1139/a06-005

Baker, N., Knudsen, T., Williams, A., 2017. Abstract Sifter: a comprehensive front-end system to PubMed. F1000Research 6, 2164. 10.12688/f1000research.12865.1

Barron, M.G., Vivian, D.N., Heintz, R.A., Yim, U.H., 2020. Long-Term Ecological Impacts from Oil Spills: Comparison of Exxon Valdez, Hebei Spirit, and Deepwater Horizon. Environmental Science & Technology 54, 6456–6467. 10.1021/acs.est.9b05020

Baudiffier, D., Audouze, K., Armant, O., Frelon, S., Charles, S., Beaudouin, R., Cosio, C., Payrastre, L., Siaussat, D., Burgeot, T., Mauffret, A., Degli Esposti, D., Mougin, C., Delaunay, D., Coumoul, X., 2024. Editorial trend: adverse outcome pathway (AOP) and computational strategy — towards new perspectives in ecotoxicology. Environmental Science and Pollution Research 31, 6587–6596. 10.1007/s11356-023-30647-w

Begum, M., Naik, S., Pradhan, U.K., Ezhilarasan, P., Sura, A.N., Bharathi, M.D., Muthukumar, C., Karthikeyan, P., Iyyappan, M., Gopinath, G., Dash, S.K., Usha, T., Panda, U.S., Mishra, P., Murthy, M.V.R., 2022. Assessment of Ennore Oil Spill 2017 on Chennai Coastal Water and Biota, in: OCEANS 2022 - Chennai. Presented at the OCEANS 2022 - Chennai, IEEE, pp. 1–7. 10.1109/OCEANSChennai45887.2022.9775218

Belanger, S., Barron, M., Craig, P., Dyer, S., Galay-Burgos, M., Hamer, M., Marshall, S., Posthuma, L., Raimondo, S., Whitehouse, P., 2017. Future needs and recommendations in the development of species sensitivity distributions: Estimating toxicity thresholds for aquatic ecological communities and assessing impacts of chemical exposures. Integrated Environmental Assessment and Management 13, 664– 674. 10.1002/ieam.1841

Beyer, J., Trannum, H.C., Bakke, T., Hodson, P.V., Collier, T.K., 2016. Environmental effects of the Deepwater Horizon oil spill: A review. Marine Pollution Bulletin 110, 28–51. 10.1016/j.marpolbul.2016.06.027

Bierkens, J., Geerts, L., 2014. Environmental hazard and risk characterisation of petroleum substances: A guided “walking tour” of petroleum hydrocarbons. Environment International 66, 182–193. 10.1016/j.envint.2014.01.030

Booc, F., Thornton, C., Lister, A., MacLatchy, D., Willett, K.L., 2014. Benzo[a]pyrene Effects on Reproductive Endpoints in Fundulus heteroclitus. Toxicological Sciences 140, 73–82. 10.1093/toxsci/kfu064

Cavanaugh, J.E., Neath, A.A., 2019. The Akaike information criterion: Background, derivation, properties, application, interpretation, and refinements. WIREs Computational Statistics 11, e1460. 10.1002/wics.1460

Chai, Z., Zhao, C., Jin, Y., Wang, Y., Zou, P., Ling, X., Yang, H., Zhou, N., Chen, Q., Sun, L., Chen, W., Ao, L., Cao, J., Liu, J., 2021. Generating adverse outcome pathway (AOP) of inorganic arsenic-induced adult male reproductive impairment via integration of phenotypic analysis in comparative toxicogenomics database (CTD) and AOP wiki. Toxicology and Applied Pharmacology 411, 115370. 10.1016/j.taap.2020.115370

Chen, J., Fan, B., Li, J., Wang, X., Li, W., Cui, L., Liu, Z., 2020. Development of human health ambient water quality criteria of 12 polycyclic aromatic hydrocarbons (PAH) and risk assessment in China. Chemosphere 252, 126590. 10.1016/j.chemosphere.2020.126590

Corrales, J., Fang, X., Thornton, C., Mei, W., Barbazuk, W.B., Duke, M., Scheffler, B.E., Willett, K.L., 2014. Effects on specific promoter DNA methylation in zebrafish embryos and larvae following benzo[a]pyrene exposure. Comparative Biochemistry and Physiology Part C: Toxicology & Pharmacology 163, 37–46. 10.1016/j.cbpc.2014.02.005

Davis, A.P., Wiegers, T.C., Johnson, R.J., Sciaky, D., Wiegers, J., Mattingly, C.J., 2023. Comparative Toxicogenomics Database (CTD): update 2023. Nucleic Acids Research 51, D1257–D1262. 10.1093/nar/gkac833

del Giudice, G., Serra, A., Pavel, A., Torres Maia, M., Saarimäki, L.A., Fratello, M., Federico, A., Alenius, H., Fadeel, B., Greco, D., 2024. A Network Toxicology Approach for Mechanistic Modelling of Nanomaterial Hazard and Adverse Outcomes. Advanced Science 2400389. 10.1002/advs.202400389

Dionisio, K.L., Phillips, K., Price, P.S., Grulke, C.M., Williams, A., Biryol, D., Hong, T., Isaacs, K.K., 2018. The Chemical and Products Database, a resource for exposure-relevant data on chemicals in consumer products. Scientific Data 5, 180125. 10.1038/sdata.2018.125

Dix, D.J., Houck, K.A., Martin, M.T., Richard, A.M., Setzer, R.W., Kavlock, R.J., 2007. The ToxCast Program for Prioritizing Toxicity Testing of Environmental Chemicals. Toxicological Sciences 95, 5–12. 10.1093/toxsci/kfl103

Djoumbou Feunang, Y., Eisner, R., Knox, C., Chepelev, L., Hastings, J., Owen, G., Fahy, E., Steinbeck, C., Subramanian, S., Bolton, E., Greiner, R., Wishart, D.S., 2016. ClassyFire: automated chemical classification with a comprehensive, computable taxonomy. Journal of Cheminformatics 8, 61. 10.1186/s13321-016-0174-y

Dowse, R., Tang, D., Palmer, C.G., Kefford, B.J., 2013. Risk assessment using the species sensitivity distribution method: Data quality versus data quantity. Environmental Toxicology and Chemistry 32, 1360–1369. 10.1002/etc.2190

Eom, I.C., Rast, C., Veber, A.M., Vasseur, P., 2007. Ecotoxicity of a polycyclic aromatic hydrocarbon (PAH)-contaminated soil. Ecotoxicology and Environmental Safety 67, 190–205. 10.1016/j.ecoenv.2006.12.020

EPA, U.S., 2002. Methods for Measuring the Acute Toxicity of Effluents and Receiving Waters to Freshwater and Marine Organisms. Fifth Edition (No. 821-R-02–012). U.S. Environmental Protection Agency.

EPA, U.S., 1998. Guidelines for Ecological Risk Assessment. U.S. Environmental Protection Agency.

EPA, U.S., 1979. Water-related environmental fate of 129 priority pollutants: Volume II: Halogenated aliphatic hydrocarbons, halogenated ethers, monocyclic aromatics, phthalate esters, polycyclic aromatic hydrocarbons, nitrosamines, and miscellaneous compounds. U.S. Environmental Protection Agency.

Etterson, M., 2020. Technical Manual: SSD Toolbox Version 1.0. U.S. Environmental Protection Agency.

Fay, K.A., Villeneuve, D.L., LaLone, C.A., Song, Y., Tollefsen, K.E., Ankley, G.T., 2017. Practical approaches to adverse outcome pathway development and weight-of-evidence evaluation as illustrated by ecotoxicological case studies. Environmental Toxicology and Chemistry 36, 1429–1449. 10.1002/etc.3770

Feshuk, M., Kolaczkowski, L., Dunham, K., Davidson-Fritz, S.E., Carstens, K.E., Brown, J., Judson, R.S., Paul Friedman, K., 2023. The ToxCast pipeline: updates to curve-fitting approaches and database structure. Frontiers in Toxicology 5, 1275980.

Fox, D.R., van Dam, R.A., Fisher, R., Batley, G.E., Tillmanns, A.R., Thorley, J., Schwarz, C.J., Spry, D.J., McTavish, K., 2021. Recent Developments in Species Sensitivity Distribution Modeling. Environmental Toxicology and Chemistry 40, 293–308. 10.1002/etc.4925

Fröhlich, H., Speer, N., Poustka, A., Beißbarth, T., 2007. GOSim – an R-package for computation of information theoretic GO similarities between terms and gene products. BMC Bioinformatics 8, 166. 10.1186/1471-2105-8-166

Gustafson, J.B., Tell, J.G., Orem, D., 1997. Selection of Representative TPH Fractions Based on Fate and Transport Considerations, Volume 3 of Total Petroleum Hydrocarbon Criteria Working Group Series. Amherst Scientific Publishers, Amherst, Massachusetts, USA.

Hagberg, A.A., Schult, D.A., Swart, P.J., 2008. Exploring Network Structure, Dynamics, and Function using NetworkX, in: Varoquaux, G., Vaught, T., Millman, J. (Eds.), Proceedings of the 7th Python in Science Conference (SciPy 2008). Presented at the SciPy 2008, pp. 11–15.

Honda, M., Suzuki, N., 2020. Toxicities of Polycyclic Aromatic Hydrocarbons for Aquatic Animals. International Journal of Environmental Research and Public Health 17, 1363. 10.3390/ijerph17041363

Hussar, E., Richards, S., Lin, Z.-Q., Dixon, R.P., Johnson, K.A., 2012. Human Health Risk Assessment of 16 Priority Polycyclic Aromatic Hydrocarbons in Soils of Chattanooga, Tennessee, USA. Water, Air, & Soil Pollution 223, 5535–5548. 10.1007/s11270-012-1265-7

Jagiello, K., Judzinska, B., Sosnowska, A., Lynch, I., Halappanavar, S., Puzyn, T., 2022. Using AOP-Wiki to support the ecotoxicological risk assessment of nanomaterials: first steps in the development of novel adverse outcome pathways. Environmental Science: Nano 9, 1675–1684. 10.1039/D1EN01127H

Jaylet, T., Coustillet, T., Jornod, F., Margaritte-Jeannin, P., Audouze, K., 2023. AOP-helpFinder 2.0: Integration of an event-event searches module. Environment International 177, 108017. 10.1016/j.envint.2023.108017

Jeong, J., Choi, J., 2019. Adverse outcome pathways potentially related to hazard identification of microplastics based on toxicity mechanisms. Chemosphere 231, 249–255. 10.1016/j.chemosphere.2019.05.003

Jornod, F., Jaylet, T., Blaha, L., Sarigiannis, D., Tamisier, L., Audouze, K., 2022. AOP-helpFinder webserver: a tool for comprehensive analysis of the literature to support adverse outcome pathways development. Bioinformatics 38, 1173–1175. 10.1093/bioinformatics/btab750

Judson, R., Houck, K., Martin, M., Richard, A.M., Knudsen, T.B., Shah, I., Little, S., Wambaugh, J., Woodrow Setzer, R., Kothya, P., Phuong, J., Filer, D., Smith, D., Reif, D., Rotroff, D., Kleinstreuer, N., Sipes, N., Xia, M., Huang, R., Crofton, K., Thomas, R.S., 2016. Analysis of the Effects of Cell Stress and Cytotoxicity on In Vitro Assay Activity Across a Diverse Chemical and Assay Space. Toxicological Sciences 152, 323–339. 10.1093/toxsci/kfw092

Kamo, M., 2023. Species Sensitivity Distribution in Ecological Risk Assessment, in: Kamo, M. (Ed.), Theories in Ecological Risk Assessment. Springer Nature Singapore, Singapore, pp. 103–134. 10.1007/978-981-99-0309-2_5

Karthikeyan, P., Marigoudar, Shambanagouda. R., Mohan, D., Sharma, K.V., Ramana Murthy, M.V., 2021. Prescribing sea water quality criteria for arsenic, cadmium and lead through species sensitivity distribution. Ecotoxicology and Environmental Safety 208, 111612. 10.1016/j.ecoenv.2020.111612

Keith, L.H., 2015. The Source of U.S. EPA’s Sixteen PAH Priority Pollutants. Polycyclic Aromatic Compounds 35, 147–160. 10.1080/10406638.2014.892886

Knapen, D., Angrish, M.M., Fortin, M.C., Katsiadaki, I., Leonard, M., Margiotta-Casaluci, L., Munn, S., O’Brien, J.M., Pollesch, N., Smith, L.C., Zhang, X., Villeneuve, D.L., 2018. Adverse outcome pathway networks I: Development and applications. Environmental Toxicology and Chemistry 37, 1723–1733. 10.1002/etc.4125

Kooijman, S.A.L.M., 1987. A safety factor for LC50 values allowing for differences in sensitivity among species. Water Research 21, 269–276. 10.1016/0043-1354(87)90205-3

Kramer, V.J., Etterson, M.A., Hecker, M., Murphy, C.A., Roesijadi, G., Spade, D.J., Spromberg, J.A., Wang, M., Ankley, G.T., 2011. Adverse outcome pathways and ecological risk assessment: Bridging to population-level effects. Environmental Toxicology and Chemistry 30, 64–76. 10.1002/etc.375

Kuppusamy, S., Maddela, N.R., Megharaj, M., Venkateswarlu, K., 2020. An Overview of Total Petroleum Hydrocarbons, in: Kuppusamy, S., Maddela, N.R., Megharaj, M., Venkateswarlu, K. (Eds.), Total Petroleum Hydrocarbons: Environmental Fate, Toxicity, and Remediation. Springer International Publishing, Cham, pp. 1–27. 10.1007/978-3-030-24035-6_1

Lawal, A.T., 2017. Polycyclic aromatic hydrocarbons. A review. Cogent Environmental Science 3, 1339841. 10.1080/23311843.2017.1339841

Leist, M., Ghallab, A., Graepel, R., Marchan, R., Hassan, R., Bennekou, S.H., Limonciel, A., Vinken, M., Schildknecht, S., Waldmann, T., Danen, E., van Ravenzwaay, B., Kamp, H., Gardner, I., Godoy, P., Bois, F.Y., Braeuning, A., Reif, R., Oesch, F., Drasdo, D., Höhme, S., Schwarz, M., Hartung, T., Braunbeck, T., Beltman, J., Vrieling, H., Sanz, F., Forsby, A., Gadaleta, D., Fisher, C., Kelm, J., Fluri, D., Ecker, G., Zdrazil, B., Terron, A., Jennings, P., van der Burg, B., Dooley, S., Meijer, A.H., Willighagen, E., Martens, M., Evelo, C., Mombelli, E., Taboureau, O., Mantovani, A., Hardy, B., Koch, B., Escher, S., van Thriel, C., Cadenas, C., Kroese, D., van de Water, B., Hengstler, J.G., 2017. Adverse outcome pathways: opportunities, limitations and open questions. Archives of Toxicology 91, 3477–3505. 10.1007/s00204-017-2045-3

Lin, S., Ren, A., Wang, L., Huang, Y., Wang, Y., Wang, C., Greene, N.D., 2018. Oxidative Stress and Apoptosis in Benzo[a]pyrene-Induced Neural Tube Defects. Free Radical Biology and Medicine 116, 149–158. 10.1016/j.freeradbiomed.2018.01.004

Logeshwaran, P., Megharaj, M., Chadalavada, S., Bowman, M., Naidu, R., 2018. Petroleum hydrocarbons (PH) in groundwater aquifers: An overview of environmental fate, toxicity, microbial degradation and risk-based remediation approaches. Environmental Technology & Innovation 10, 175–193. 10.1016/j.eti.2018.02.001

Machin, D., Bryant, T., Altman, D., Gardner, M. (Eds.), 2000. Statistics with Confidence: Confidence Intervals and Statistical Guidelines, 2nd ed. BMJ Books.

Malott, K.F., Leon Parada, K., Lee, M., Swanson, E., Luderer, U., 2022. Gestational Benzo[a]pyrene Exposure Destroys F1 Ovarian Germ Cells Through Mitochondrial Apoptosis Pathway and Diminishes Surviving Oocyte Quality. Toxicological Sciences 190, 23–40. 10.1093/toxsci/kfac086

Méndez-Fernández, L., Casado-Martínez, C., Martínez-Madrid, M., Moreno-Ocio, I., Costas, N., Pardo, I., Rodriguez, P., 2019. Derivation of sediment Hg quality standards based on ecological assessment in river basins. Environmental Pollution 245, 1000–1013. 10.1016/j.envpol.2018.11.068

National Research Council, 2007. Toxicity Testing in the 21st Century: A Vision and a Strategy. The National Academies Press, Washington, DC. 10.17226/11970

Olker, J.H., Elonen, C.M., Pilli, A., Anderson, A., Kinziger, B., Erickson, S., Skopinski, M., Pomplun, A., LaLone, C.A., Russom, C.L., Hoff, D., 2022. The ECOTOXicology Knowledgebase: A Curated Database of Ecologically Relevant Toxicity Tests to Support Environmental Research and Risk Assessment. Environmental Toxicology and Chemistry 41, 1520–1539. 10.1002/etc.5324

Pandelides, Z., Sturgis, M.C., Thornton, C., Aluru, N., Willett, K.L., 2023. Benzo[a]pyrene-induced multigenerational changes in gene expression, behavior, and DNA methylation are primarily influenced by paternal exposure. Toxicology and Applied Pharmacology 469, 116545. 10.1016/j.taap.2023.116545

Pasparakis, C., Esbaugh, A.J., Burggren, W., Grosell, M., 2019. Physiological impacts of Deepwater Horizon oil on fish. Comparative Biochemistry and Physiology Part C: Toxicology & Pharmacology 224, 108558. 10.1016/j.cbpc.2019.06.002

Posthuma, L., Suter II, G.W., Traas, T.P. (Eds.), 2001. Species Sensitivity Distributions in Ecotoxicology. CRC Press, Boca Raton. 10.1201/9781420032314

Posthuma, L., van Gils, J., Zijp, M.C., van de Meent, D., de Zwart, D., 2019. Species sensitivity distributions for use in environmental protection, assessment, and management of aquatic ecosystems for 12_386 chemicals. Environmental Toxicology and Chemistry 38, 905–917. 10.1002/etc.4373

Potter, T.L., Simmons, K.E., 1998. Composition of Petroleum Mixtures, Volume 2 of Total Petroleum Hydrocarbon Criteria Working Group Series. Amherst Scientific Publishers, Amherst, Massachusetts, USA.

Pritsos, K.L., Perez, C.R., Muthumalage, T., Dean, K.M., Cacela, D., Hanson-Dorr, K., Cunningham, F., Bursian, S.J., Link, J.E., Shriner, S., Horak, K., Pritsos, C.A., 2017. Dietary intake of Deepwater Horizon oil-injected live food fish by double-crested cormorants resulted in oxidative stress. Ecotoxicology and Environmental Safety 146, 62–67. 10.1016/j.ecoenv.2017.06.067

Ravichandran, J., Karthikeyan, B.S., Samal, A., 2022. Investigation of a derived adverse outcome pathway (AOP) network for endocrine-mediated perturbations. Science of The Total Environment 826, 154112. 10.1016/j.scitotenv.2022.154112

Russom, C.L., LaLone, C.A., Villeneuve, D.L., Ankley, G.T., 2014. Development of an adverse outcome pathway for acetylcholinesterase inhibition leading to acute mortality. Environmental Toxicology and Chemistry 33, 2157–2169. 10.1002/etc.2662

Sahoo, A.K., Chivukula, N., Madgaonkar, S.R., Ramesh, K., Marigoudar, S.R., Sharma, K.V., Samal, A., name2024a. Leveraging integrative toxicogenomic approach towards development of stressor-centric adverse outcome pathway networks for plastic additives. bioRxiv 2024.03.27.586984.

Sahoo, A.K., Chivukula, N., Ramesh, K., Singha, J., Marigoudar, S.R., Sharma, K.V., Samal, A., 2024b. An integrative data-centric approach to derivation and characterization of an adverse outcome pathway network for cadmium-induced toxicity. Science of The Total Environment 920, 170968. 10.1016/j.scitotenv.2024.170968

Sammarco, P.W., Kolian, S.R., Warby, R.A.F., Bouldin, J.L., Subra, W.A., Porter, S.A., 2013. Distribution and concentrations of petroleum hydrocarbons associated with the BP/Deepwater Horizon Oil Spill, Gulf of Mexico. Marine Pollution Bulletin 73, 129–143. 10.1016/j.marpolbul.2013.05.029

Scholz, S., Sela, E., Blaha, L., Braunbeck, T., Galay-Burgos, M., García-Franco, M., Guinea, J., Klüver, N., Schirmer, K., Tanneberger, K., Tobor-Kapłon, M., Witters, H., Belanger, S., Benfenati, E., Creton, S., Cronin, M.T.D., Eggen, R.I.L., Embry, M., Ekman, D., Gourmelon, A., Halder, M., Hardy, B., Hartung, T., Hubesch, B., Jungmann, D., Lampi, M.A., Lee, L., Léonard, M., Küster, E., Lillicrap, A., Luckenbach, T., Murk, A.J., Navas, J.M., Peijnenburg, W., Repetto, G., Salinas, E., Schüürmann, G., Spielmann, H., Tollefsen, K.E., Walter-Rohde, S., Whale, G., Wheeler, J.R., Winter, M.J., 2013. A European perspective on alternatives to animal testing for environmental hazard identification and risk assessment. Regulatory Toxicology and Pharmacology 67, 506–530. 10.1016/j.yrtph.2013.10.003

Schwarz, C., Tillmanns, A., 2019. Improving statistical methods to derive species sensitivity distribution (No. WSS2019-0), Water Science Series. Province of British Columbia, Victoria.

Shannon, P., Markiel, A., Ozier, O., Baliga, N.S., Wang, J.T., Ramage, D., Amin, N., Schwikowski, B., Ideker, T., 2003. Cytoscape: A Software Environment for Integrated Models of Biomolecular Interaction Networks. Genome Research 13, 2498–2504. 10.1101/gr.1239303

Sharma, H., Jain, V.K., Khan, Z.H., 2007. Characterization and source identification of polycyclic aromatic hydrocarbons (PAHs) in the urban environment of Delhi. Chemosphere 66, 302–310. 10.1016/j.chemosphere.2006.05.003

Sivagami, K., Jaa Vignesh, V., Tamizhdurai, P., Rajasekhar, B., Sakthipriya, N., Nambi, I.M., 2019. Studies on short term weathering of spilled oil along Chennai coast in South India. Journal of Cleaner Production 230, 1410–1420. 10.1016/j.jclepro.2019.05.119

Sopian, N.A., Jalaludin, J., Abu Bakar, S., Hamedon, T.R., Latif, M.T., 2021. Exposure to Particulate PAHs on Potential Genotoxicity and Cancer Risk among School Children Living Near the Petrochemical Industry. International Journal of Environmental Research and Public Health 18, 2575. 10.3390/ijerph18052575

Sui, L., Nie, J., Xiao, P., Yan, K., Zhang, Huiting, Liu, J., Zhang, Hengye, Cui, K., Lu, K., Liang, X., 2020. Maternal benzo[a]pyrene exposure is correlated with the meiotic arrest and quality deterioration of offspring oocytes in mice. Reproductive Toxicology 93, 10–18. 10.1016/j.reprotox.2019.12.003

Takes, F.W., Kosters, W.A., 2011. Determining the Diameter of Small World Networks, in: Proceedings of the 20th ACM International Conference on Information and Knowledge Management, CIKM‑11. Association for Computing Machinery, New York, NY, USA, pp. 1191–1196. 10.1145/2063576.2063748

Takeshita, R., Bursian, S.J., Colegrove, K.M., Collier, T.K., Deak, K., Dean, K.M., De Guise, S., DiPinto, L.M., Elferink, C.J., Esbaugh, A.J., Griffitt, R.J., Grosell, M., Harr, K.E., Incardona, J.P., Kwok, R.K., Lipton, J., Mitchelmore, C.L., Morris, J.M., Peters, E.S., Roberts, A.P., Rowles, T.K., Rusiecki, J.A., Schwacke, L.H., Smith, C.R., Wetzel, D.L., Ziccardi, M.H., Hall, A.J., 2021. A review of the toxicology of oil in vertebrates: what we have learned following the Deepwater Horizon oil spill. Journal of Toxicology and Environmental Health, Part B 24, 355–394. 10.1080/10937404.2021.1975182

Thomas, R.S., Bahadori, T., Buckley, T.J., Cowden, J., Deisenroth, C., Dionisio, K.L., Frithsen, J.B., Grulke, C.M., Gwinn, M.R., Harrill, J.A., Higuchi, M., Houck, K.A., Hughes, M.F., Hunter, E.S., III, Isaacs, K.K., Judson, R.S., Knudsen, T.B., Lambert, J.C., Linnenbrink, M., Martin, T.M., Newton, S.R., Padilla, S., Patlewicz, G., Paul-Friedman, K., Phillips, K.A., Richard, A.M., Sams, R., Shafer, T.J., Setzer, R.W., Shah, I., Simmons, J.E., Simmons, S.O., Singh, A., Sobus, J.R., Strynar, M., Swank, A., Tornero-Valez, R., Ulrich, E.M., Villeneuve, D.L., Wambaugh, J.F., Wetmore, B.A., Williams, A.J., 2019. The Next Generation Blueprint of Computational Toxicology at the U.S. Environmental Protection Agency. Toxicological Sciences 169, 317–332. 10.1093/toxsci/kfz058

Tian, Y., Zeng, Y., Li, C., Wang, X., Liu, Q., Zhao, Y., 2020. Ecological risk assessment of petroleum hydrocarbons on aquatic organisms based on multisource data. Ecotoxicology and Environmental Safety 192, 110262. 10.1016/j.ecoenv.2020.110262

Tornero, V., Hanke, G., 2016. Chemical contaminants entering the marine environment from sea-based sources: A review with a focus on European seas. Marine Pollution Bulletin 112, 17–38. 10.1016/j.marpolbul.2016.06.091

van Straalen, N.M., Denneman, C.A.J., 1989. Ecotoxicological evaluation of soil quality criteria. Ecotoxicology and Environmental Safety 18, 241–251. 10.1016/0147-6513(89)90018-3

Villeneuve, D.L., Angrish, M.M., Fortin, M.C., Katsiadaki, I., Leonard, M., Margiotta-Casaluci, L., Munn, S., O’Brien, J.M., Pollesch, N.L., Smith, L.C., Zhang, X., Knapen, D., 2018. Adverse outcome pathway networks II: Network analytics. Environmental Toxicology and Chemistry 37, 1734–1748. 10.1002/etc.4124

Villeneuve, D.L., Crump, D., Garcia-Reyero, N., Hecker, M., Hutchinson, T.H., LaLone, C.A., Landesmann, B., Lettieri, T., Munn, S., Nepelska, M., Ottinger, M.A., Vergauwen, L., Whelan, M., 2014a. Adverse Outcome Pathway (AOP) Development I: Strategies and Principles. Toxicological Sciences 142, 312–320. 10.1093/toxsci/kfu199

Villeneuve, D.L., Crump, D., Garcia-Reyero, N., Hecker, M., Hutchinson, T.H., LaLone, C.A., Landesmann, B., Lettieri, T., Munn, S., Nepelska, M., Ottinger, M.A., Vergauwen, L., Whelan, M., 2014b. Adverse Outcome Pathway Development II: Best Practices. Toxicological Sciences 142, 321–330. 10.1093/toxsci/kfu200

Wan, T., Mo, J., Au, D.W.-T., Qin, X., Tam, N.Y.-K., Kong, R.Y.-C., Seemann, F., 2023. The role of DNA methylation on gene expression in the vertebrae of ancestrally benzo[a]pyrene exposed F1 and F3 male medaka. Epigenetics 18, 2222246. 10.1080/15592294.2023.2222246

Wang, Shiqi, Li, C., Zhang, L., Chen, Q., Wang, Shuoliang, 2024. Assessing the ecological impacts of polycyclic aromatic hydrocarbons petroleum pollutants using a network toxicity model. Environmental Research 245, 117901. 10.1016/j.envres.2023.117901

Weisman, W., 1998. Analysis of Petroleum Hydrocarbons in Environmental Media, Volume 1 of Total Petroleum Hydrocarbon Criteria Working Group Series. Amherst Scientific Publishers, Amherst, Massachusetts, USA.

Wheeler, J.R., Grist, E.P.M., Leung, K.M.Y., Morritt, D., Crane, M., 2002. Species sensitivity distributions: data and model choice. Marine Pollution Bulletin 45, 192–202. 10.1016/S0025-326X(01)00327-7

Wojciechowski, M.F., Meehan, T., 1984. Inhibition of DNA methyltransferases in vitro by benzo[a]pyrene diol epoxide-modified substrates. Journal of Biological Chemistry 259, 9711–9716. 10.1016/S0021-9258(17)42758-X

Yan, J., Wang, L., Fu, P.P., Yu, H., 2004. Photomutagenicity of 16 polycyclic aromatic hydrocarbons from the US EPA priority pollutant list. Mutation Research/Genetic Toxicology and Environmental Mutagenesis 557, 99–108. 10.1016/j.mrgentox.2003.10.004

Yanagihara, M., Hiki, K., Iwasaki, Y., 2022. Can Chemical Toxicity in Saltwater Be Predicted from Toxicity in Freshwater? A Comprehensive Evaluation Using Species Sensitivity Distributions. Environmental Toxicology and Chemistry 41, 2021–2027. 10.1002/etc.5354

Yauk, C.L., Polyzos, A., Rowan-Carroll, A., Kortubash, I., Williams, A., Kovalchuk, O., 2008. Tandem repeat mutation, global DNA methylation, and regulation of DNA methyltransferases in cultured mouse embryonic fibroblast cells chronically exposed to chemicals with different modes of action. Environmental and Molecular Mutagenesis 49, 26–35. 10.1002/em.20359

Zelinkova, Z., Wenzl, T., 2015. The Occurrence of 16 EPA PAHs in Food – A Review. Polycyclic Aromatic Compounds 35, 248–284. 10.1080/10406638.2014.918550

Zhang, Y., Tao, S., 2009. Global atmospheric emission inventory of polycyclic aromatic hydrocarbons (PAHs) for 2004. Atmospheric Environment 43, 812–819. 10.1016/j.atmosenv.2008.10.050

Zheng, G.J., Richardson, B.J., 1999. Petroleum hydrocarbons and polycyclic aromatic hydrocarbons (PAHs) in Hong Kong marine sediments. Chemosphere 38, 2625–2632. 10.1016/S0045-6535(98)00470-6

Zhuo, S., Shen, G., Zhu, Y., Du, W., Pan, X., Li, T., Han, Y., Li, B., Liu, J., Cheng, H., Xing, B., Tao, S., 2017. Source-oriented risk assessment of inhalation exposure to ambient polycyclic aromatic hydrocarbons and contributions of non-priority isomers in urban Nanjing, a megacity located in Yangtze River Delta, China. Environmental Pollution 224, 796–809. 10.1016/j.envpol.2017.01.039

