## Supplementary Figure for "Network-based investigation of petroleum hydrocarbons-induced ecotoxicological effects and their risk assessment"

**Supplementary Figures S1-S12**

**for**

**Network-based investigation of petroleum hydrocarbons-induced ecotoxicological effects and their risk assessment**

Ajaya Kumar Sahoo^a,b,1^, Shreyes Rajan Madgaonkar^a,b,1^, Nikhil Chivukula^a,b^, Panneerselvam Karthikeyan^c^, Kundhanathan Ramesh^a^, Shambanagouda Rudragouda Marigoudar^c^, Krishna Venkatarama Sharma^c^, Areejit Samal^a,b,*^

*^a^ The Institute of Mathematical Sciences (IMSc), Chennai, India*

*^b^ Homi Bhabha National Institute (HBNI), Mumbai, India*

*^c^ National Centre for Coastal Research, Ministry of Earth Sciences, Government of India, Pallikaranai, Chennai, India*

^1^A.K. Sahoo and S.R. Madgaonkar contributed equally to this work and should be considered as Joint-First authors

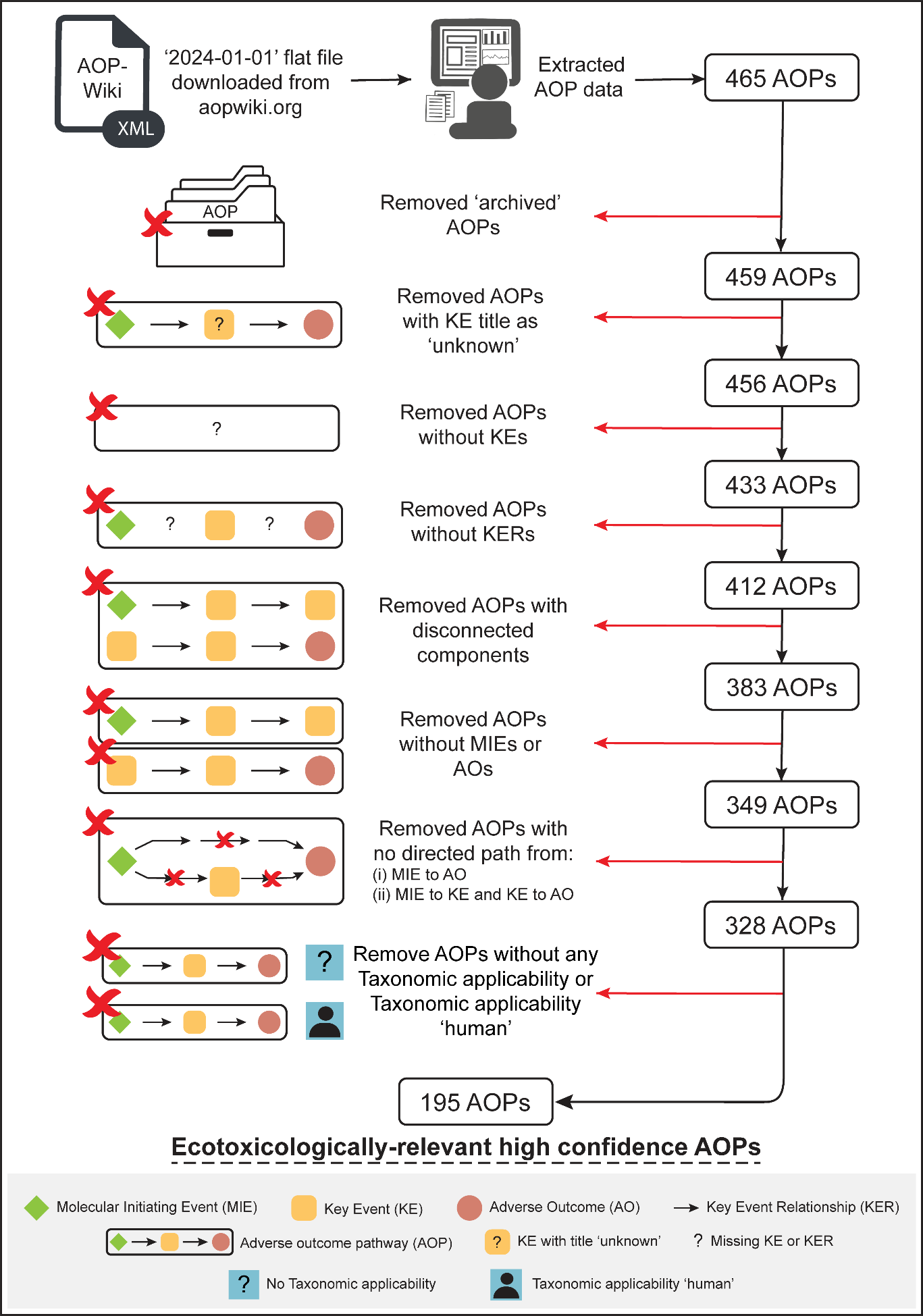

**Figure S1:** Workflow to filter ecotoxicologically-relevant high confidence adverse outcome pathways (AOPs) from AOP-Wiki by employing computation and manual curation in conjunction. (Adapted from Sahoo *et al*., 2024a).

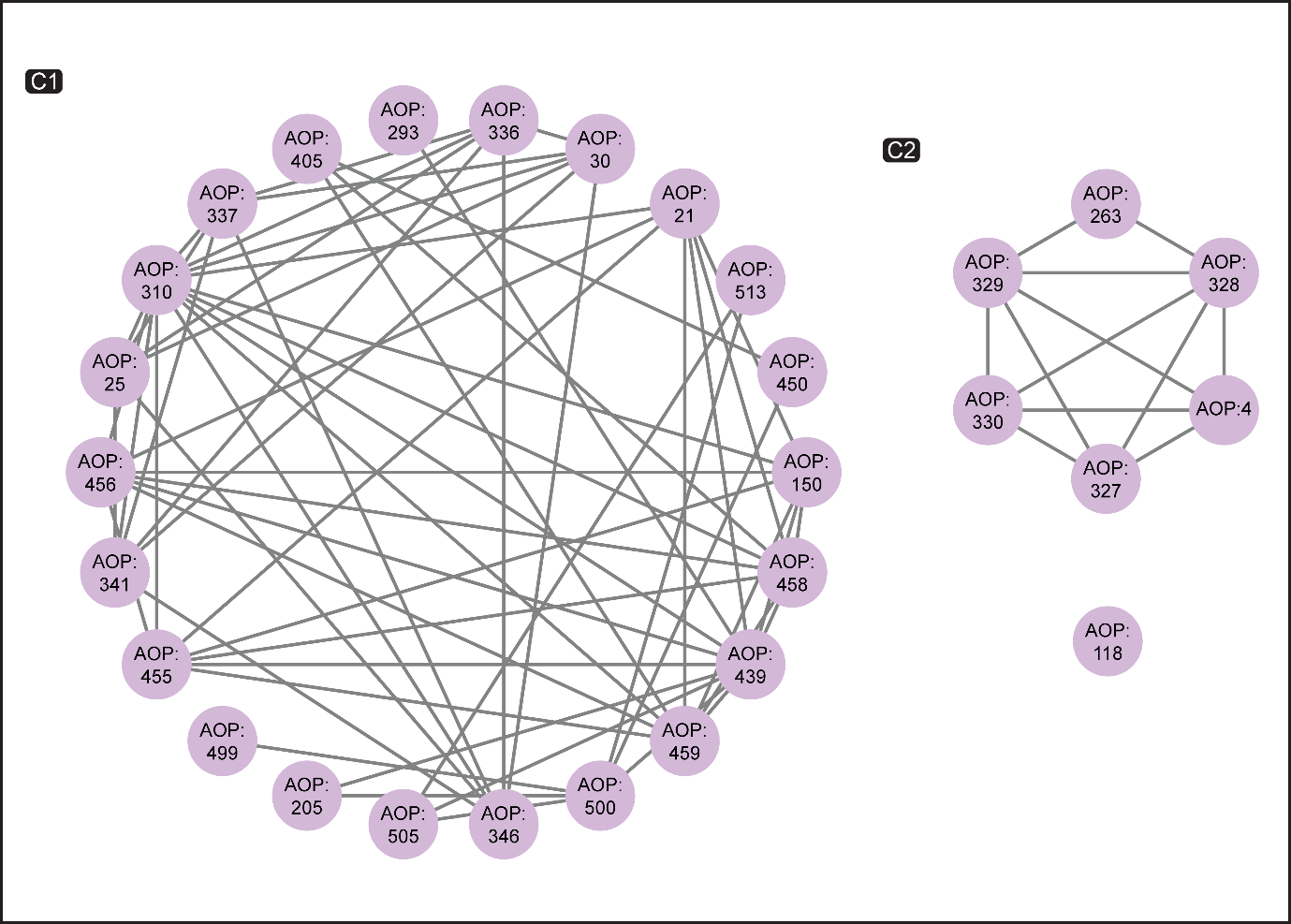

**Figure S2:** Undirected network of benzo[a]pyrene (B[a]P)-AOPs. Each node corresponds to a B[a]P-AOP and an edge between two nodes denotes that the two AOPs share at least one KE. This undirected network has 2 connected components (wherein at least two nodes are connected) which are labeled as C1, C2, and one isolated node.

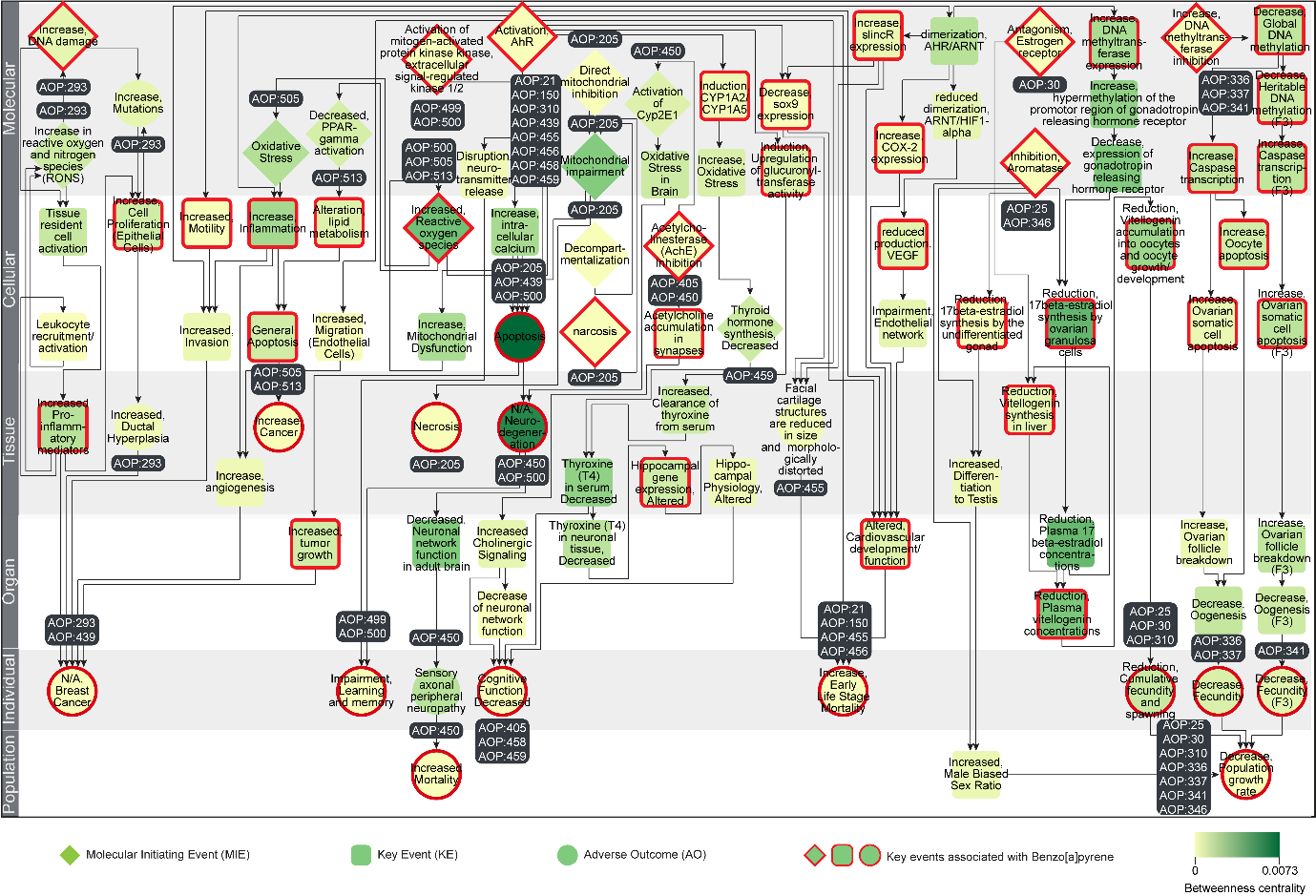

**Figure S3:** Directed network corresponding to the largest connected component (C1) in the B[a]P-AOP network, where the KEs (including MIEs and AOs) are colored based on their betweenness centrality values. The 51 KEs (including MIEs and AOs) associated with B[a]P are marked in ‘red’. In this figure, the 92 KEs are arranged vertically according to their level of biological organization.

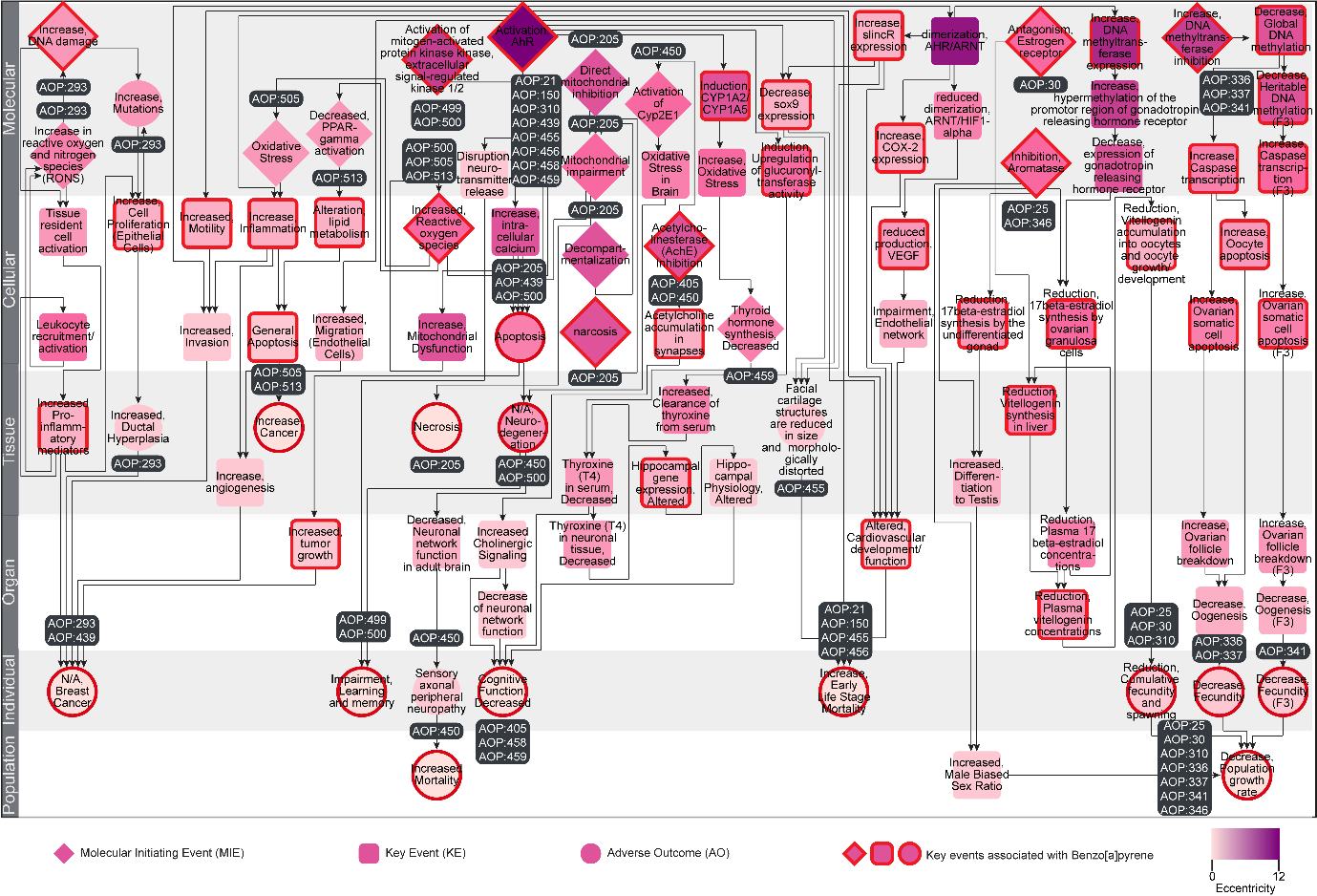

**Figure S4:** Directed network corresponding to the largest connected component (C1) in the B[a]P-AOP network, where the KEs (including MIEs and AOs) are colored based on their eccentricity values. The 51 KEs (including MIEs and AOs) associated with B[a]P are marked in ‘red’. In this figure, the 92 KEs are arranged vertically according to their level of biological organization.

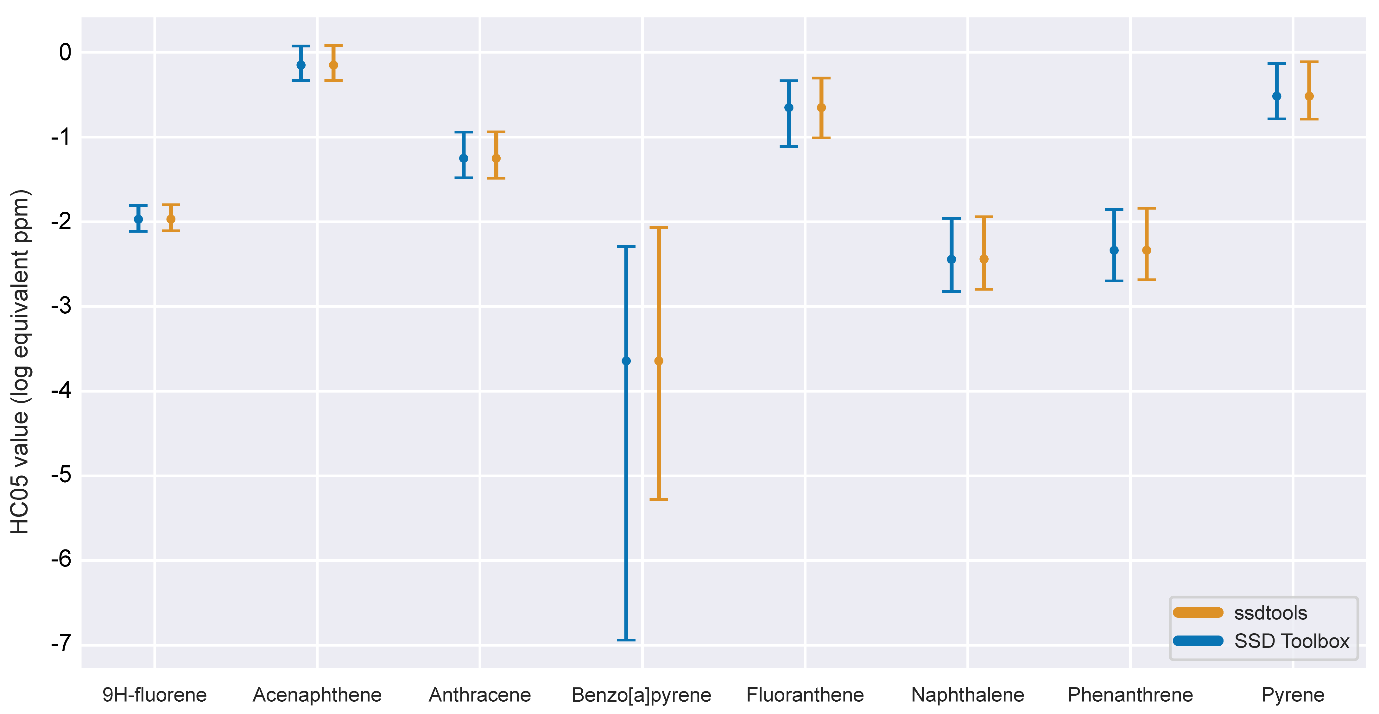

**Figure S5:** Comparison of the derived HC05 values for each of the eight priority PAHs using the corresponding best-fit model in US EPA SSD Toolbox and ssdtools.

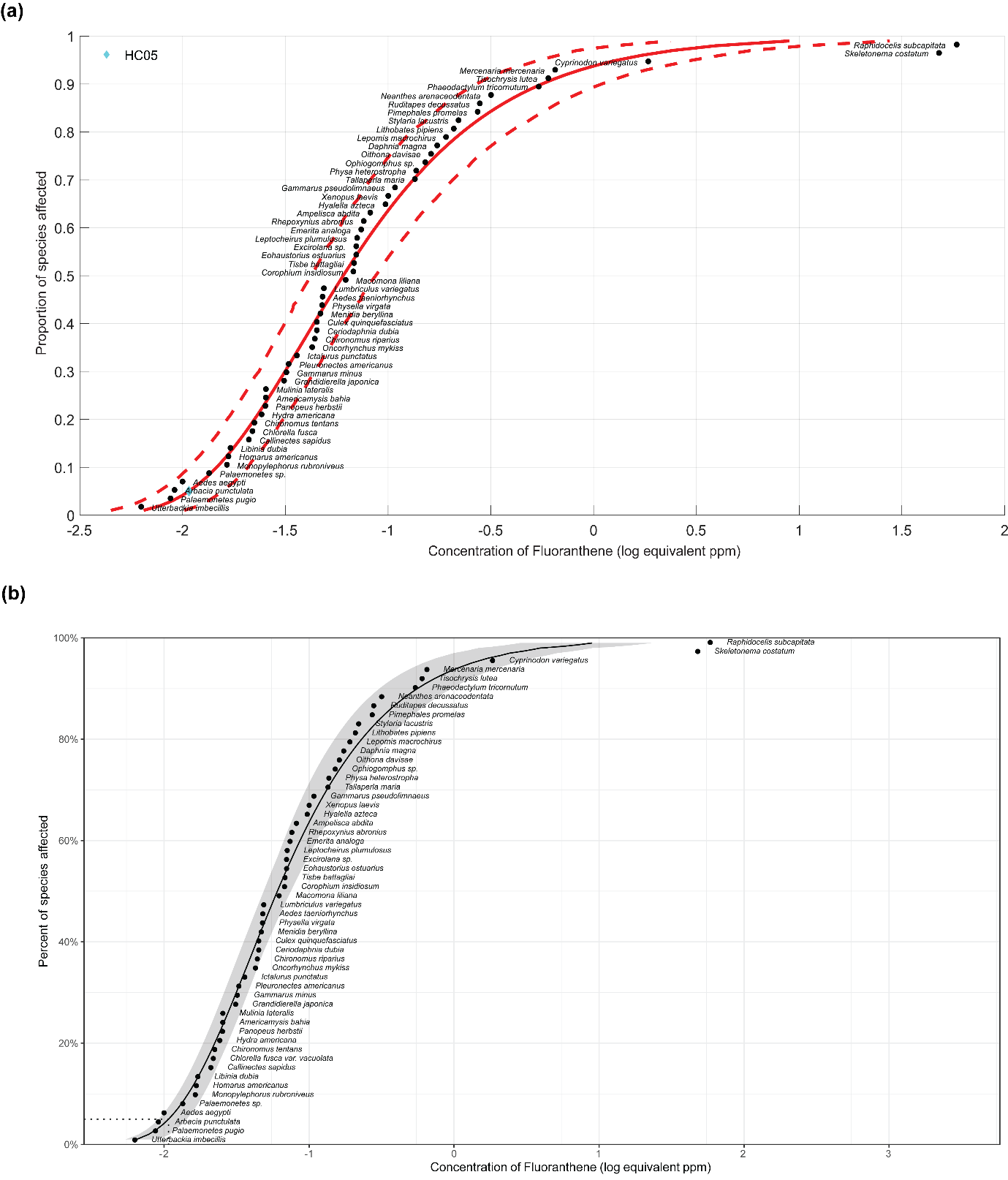

**Figure S6:** The plots of SSD for fluoranthene computed using the best-fit Log-Gumbel model.  **(a)** As determined by US EPA SSD Toolbox where the HC05 value is denoted by cyan colored diamond. **(b)** As determined by ssdtools where the HC05 value is denoted by a dotted line.

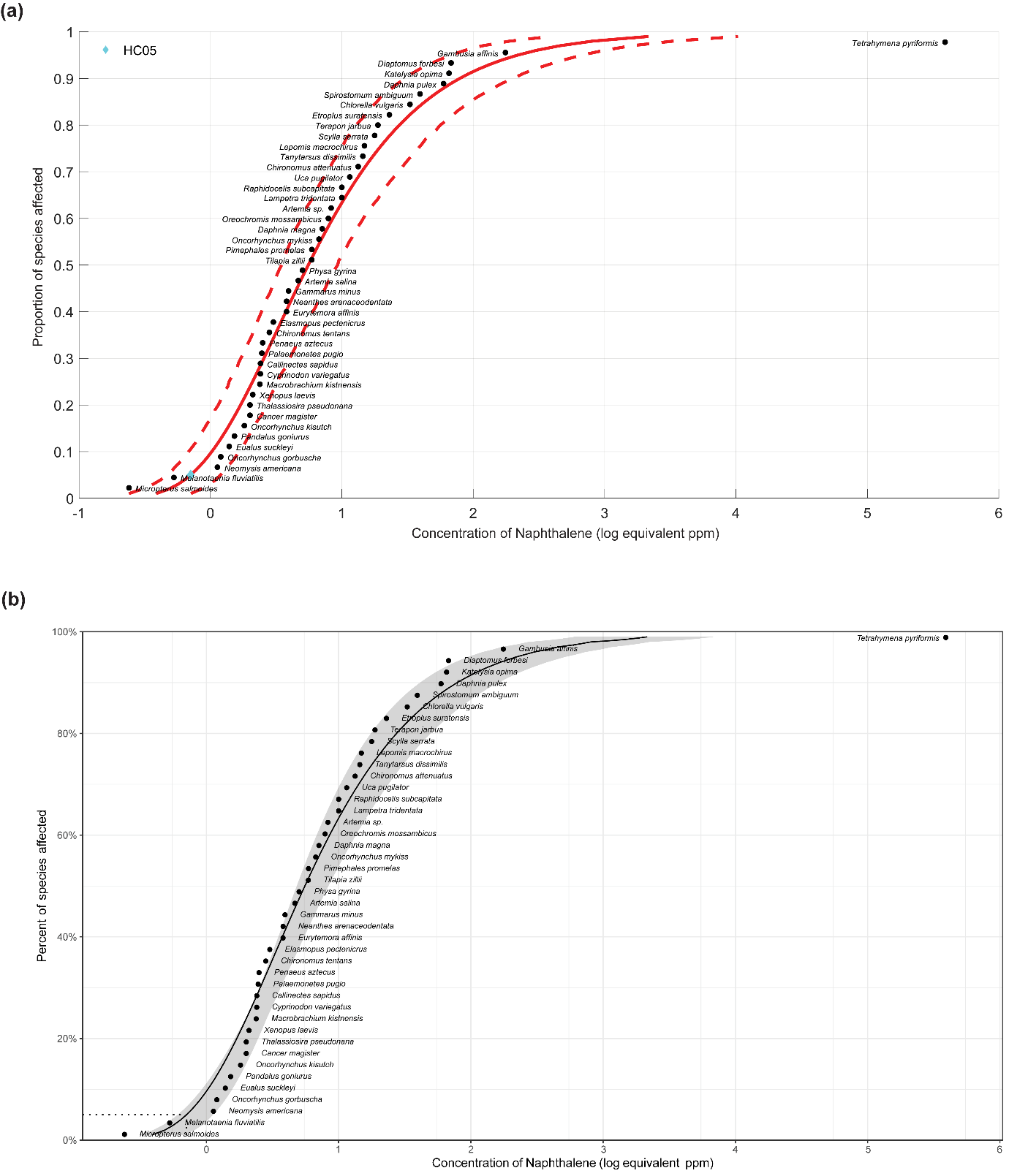

**Figure S7:** The plots of SSD for naphthalene computed using the best-fit Log-Gumbel model.  **(a)** As determined by US EPA SSD Toolbox where the HC05 value is denoted by cyan colored diamond. **(b)** As determined by ssdtools where the HC05 value is denoted by a dotted line.

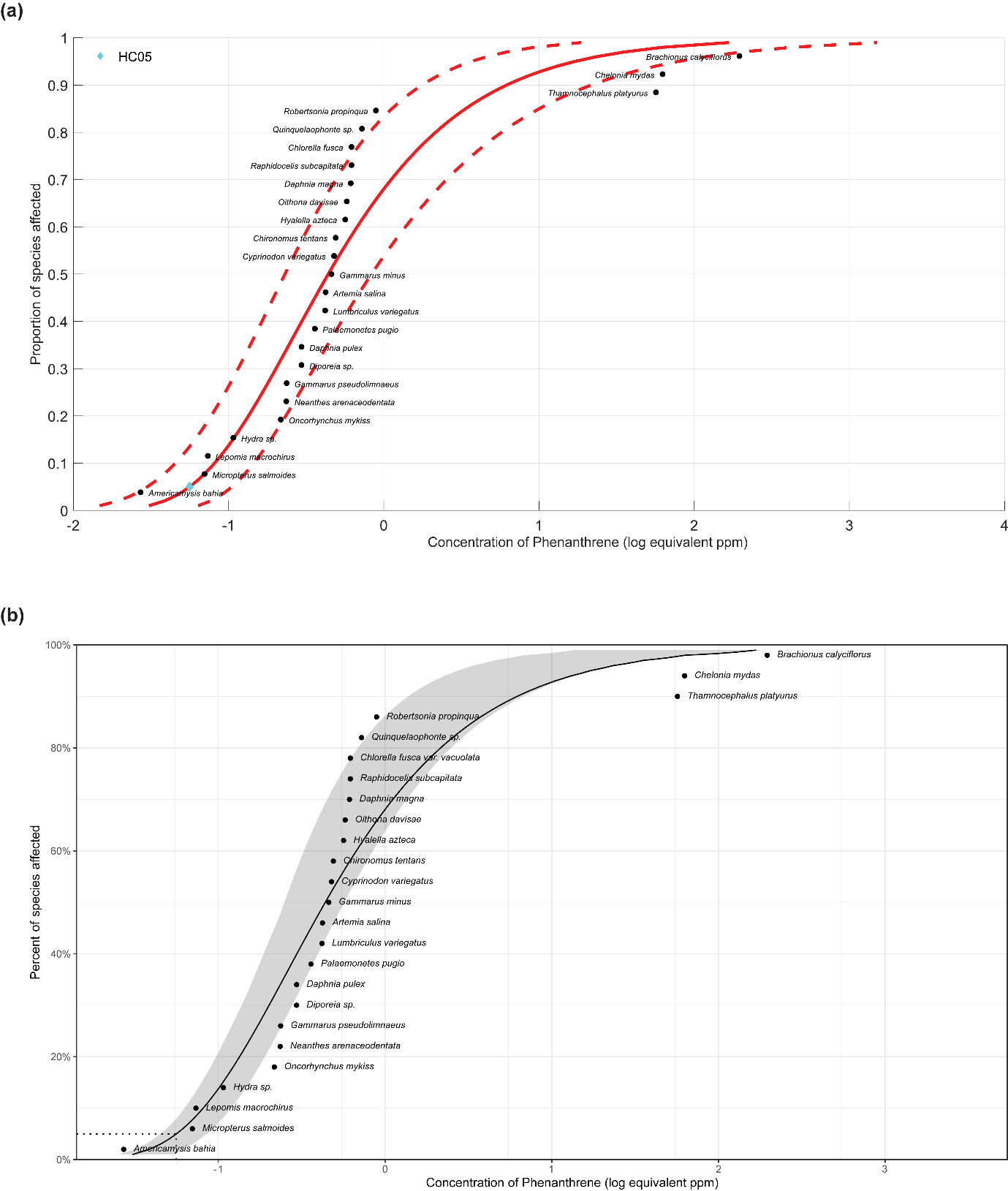

**Figure S8:** The plots of SSD for phenanthrene computed using the best-fit Log-Gumbel model.  **(a)** As determined by US EPA SSD Toolbox where the HC05 value is denoted by cyan colored diamond. **(b)** As determined by ssdtools where the HC05 value is denoted by a dotted line.

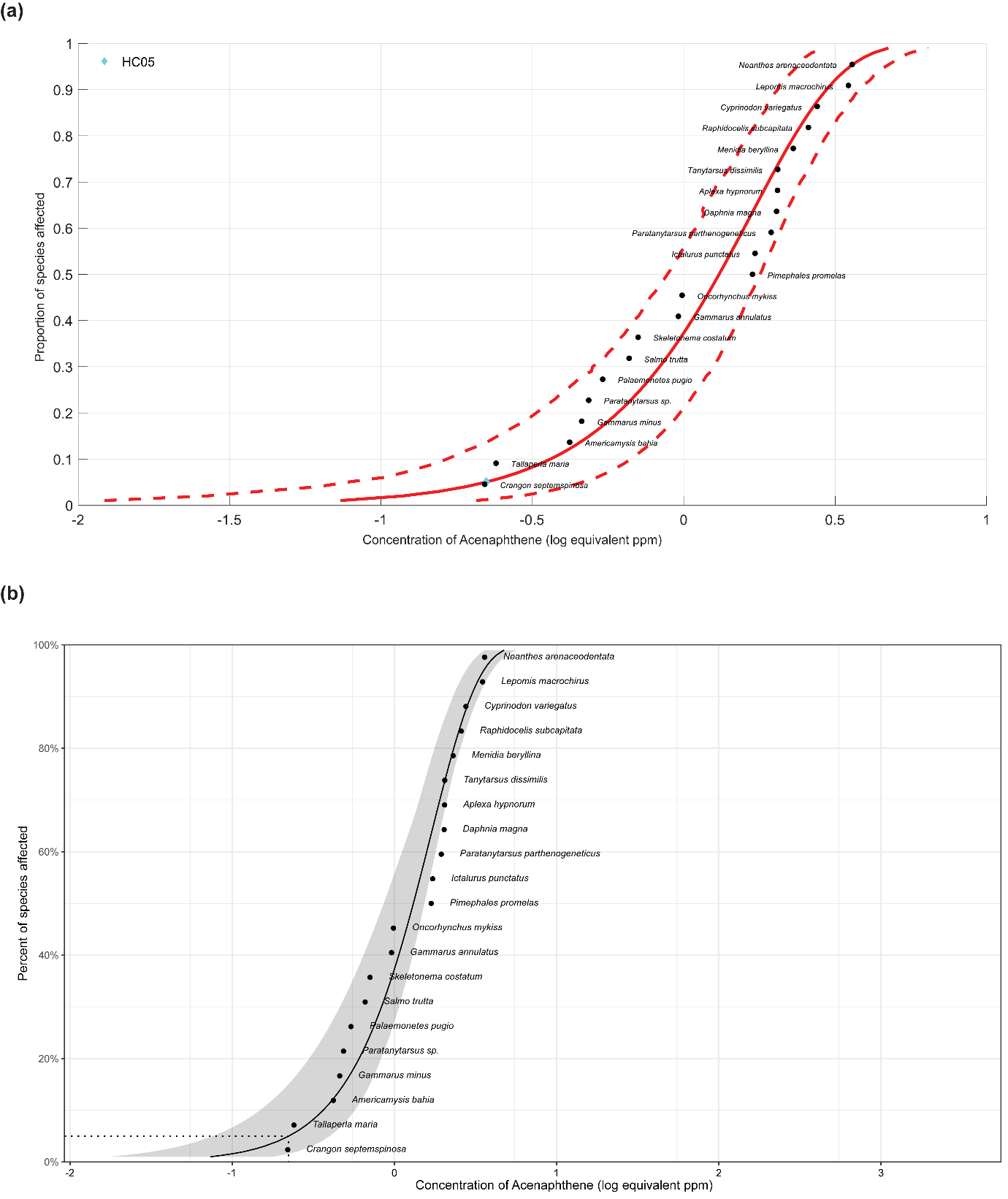

**Figure S9:** The plots of SSD for acenaphthene computed using the best-fit Weibull model.  **(a)** As determined by US EPA SSD Toolbox where the HC05 value is denoted by cyan colored diamond. **(b)** As determined by ssdtools where the HC05 value is denoted by a dotted line.

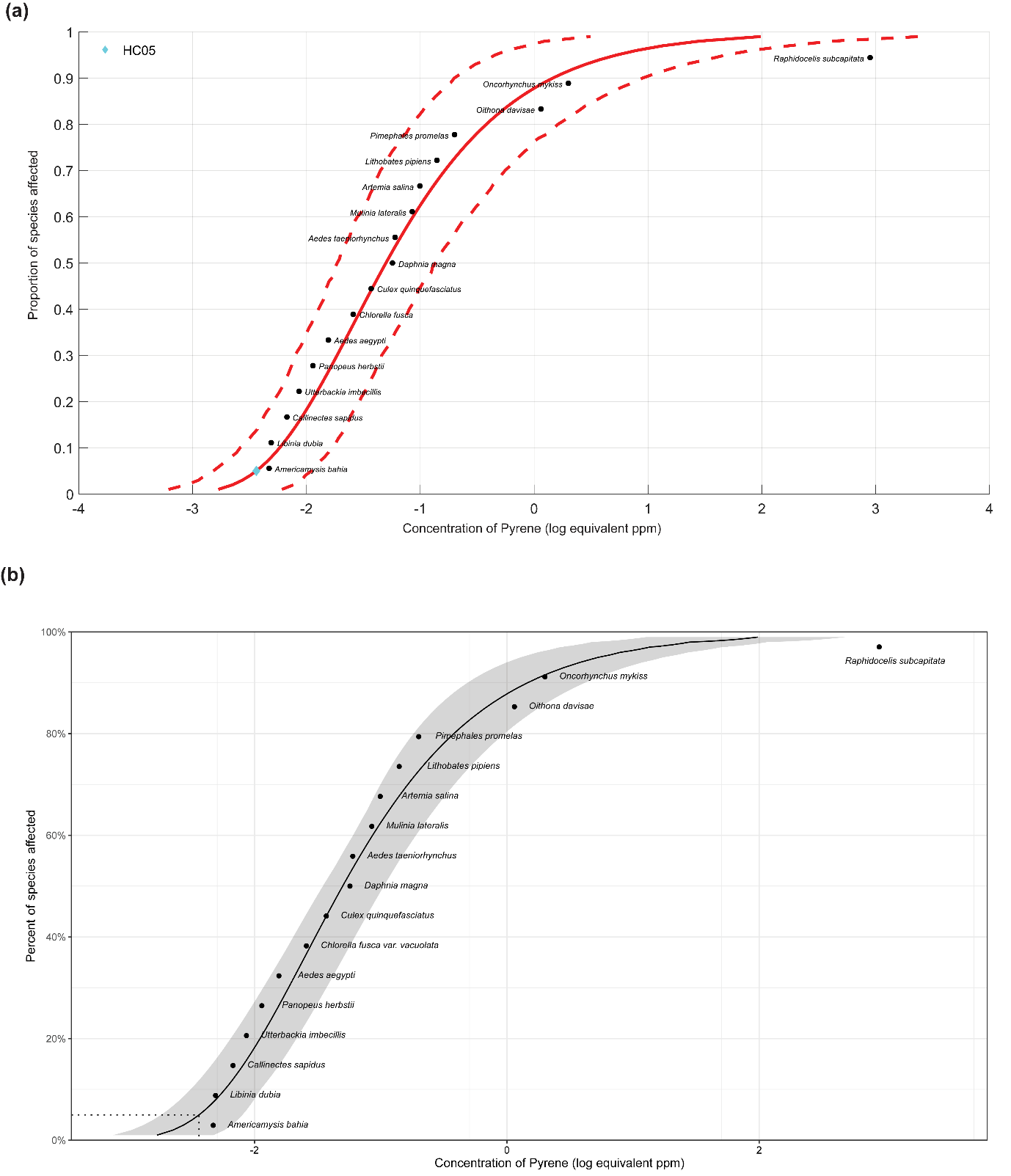

**Figure S10:** The plots of SSD for pyrene computed using the best-fit Log-Gumbel model.  **(a)** As determined by US EPA SSD Toolbox where the HC05 value is denoted by cyan colored diamond. **(b)** As determined by ssdtools where the HC05 value is denoted by a dotted line.

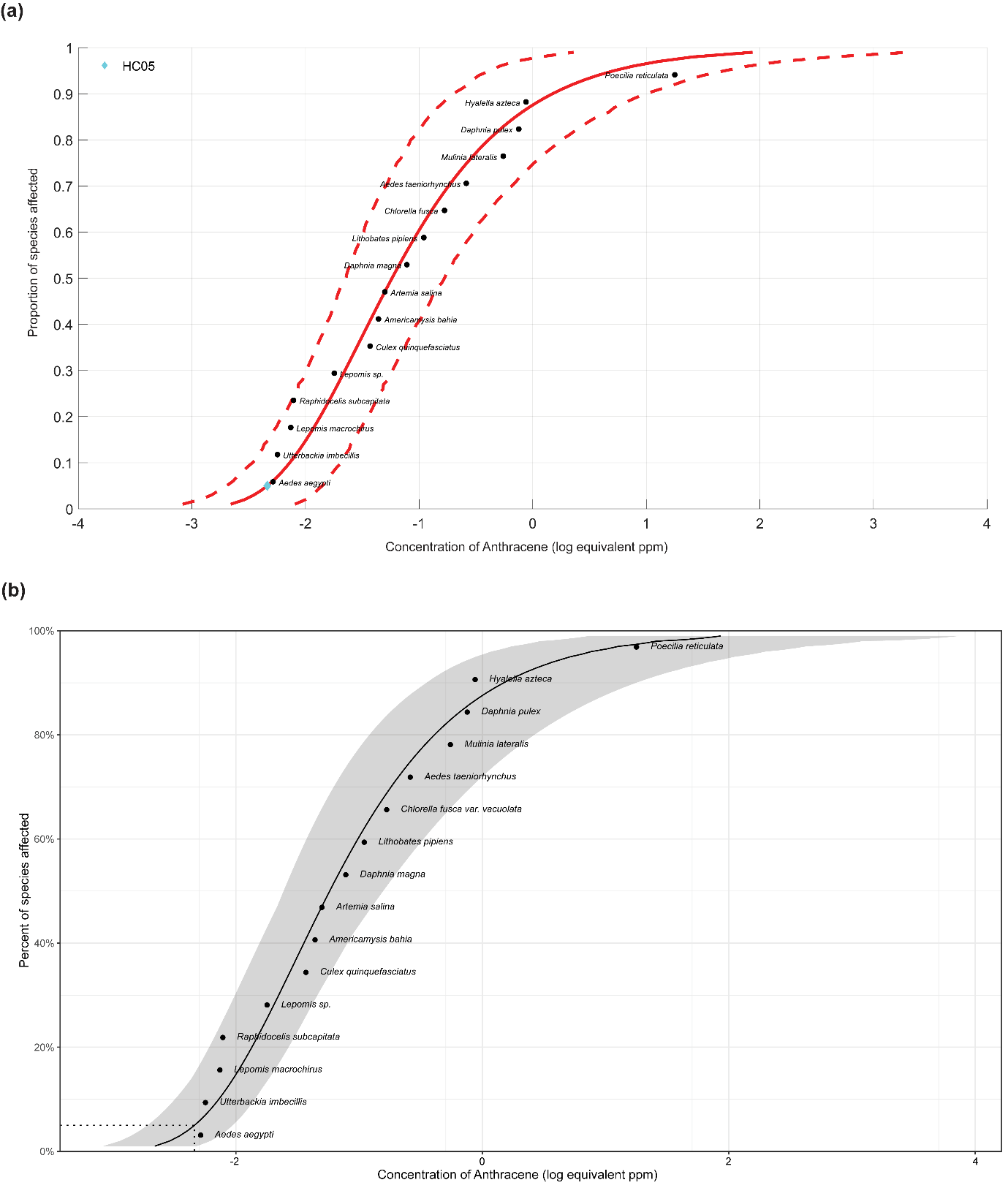

**Figure S11:** The plots of SSD for anthracene computed using the best-fit Log-Gumbel model.  **(a)** As determined by US EPA SSD Toolbox where the HC05 value is denoted by cyan colored diamond. **(b)** As determined by ssdtools where the HC05 value is denoted by a dotted line.

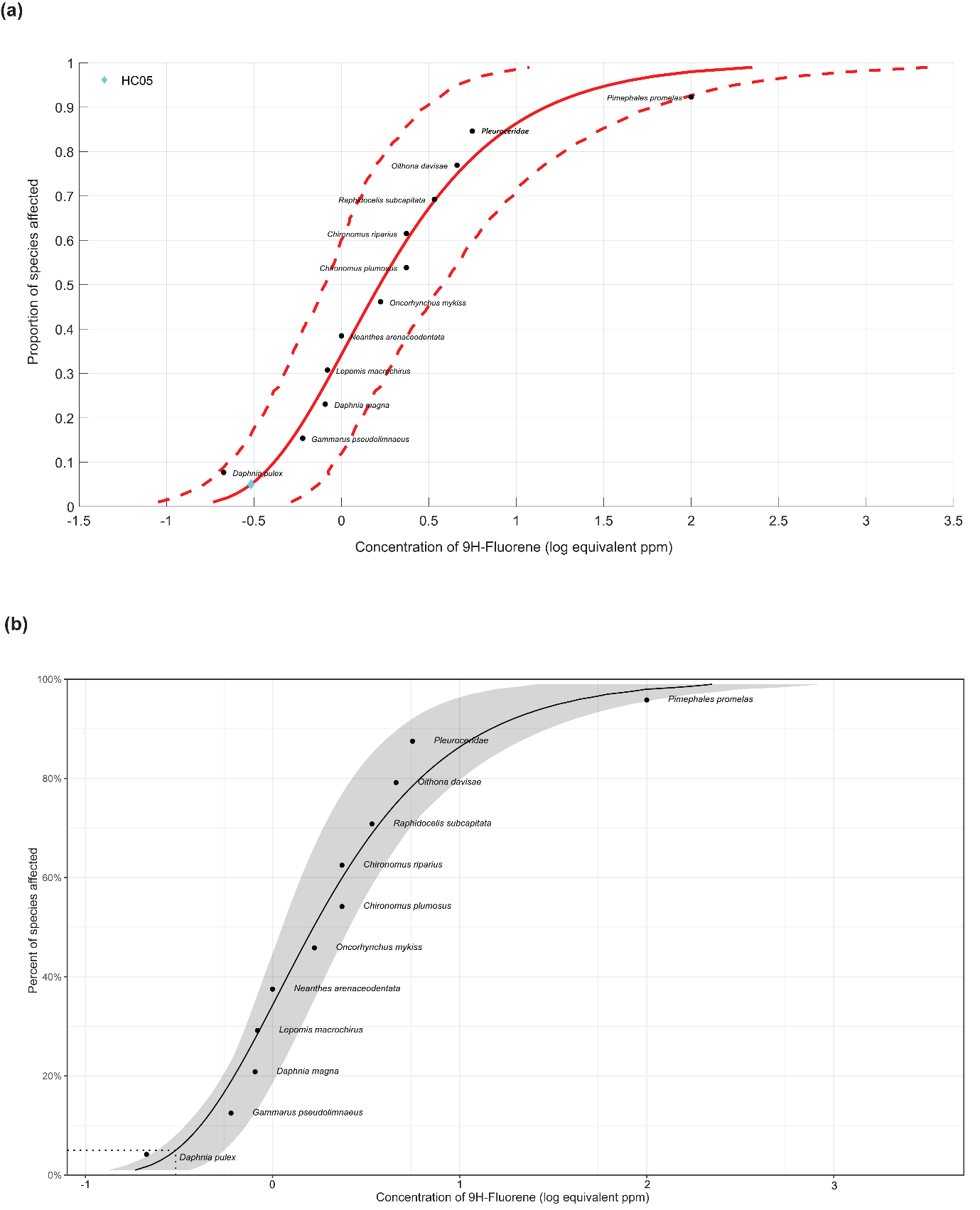

**Figure S12:** The plots of SSD for 9H-fluorene computed using the best-fit Log-Gumbel model.  **(a)** As determined by US EPA SSD Toolbox where the HC05 value is denoted by cyan colored diamond. **(b)** As determined by ssdtools where the HC05 value is denoted by a dotted line.
